# Zeb2 DNA-binding sites in neuroprogenitor cells reveal autoregulation and affirm neurodevelopmental defects, including in Mowat-Wilson Syndrome

**DOI:** 10.1101/2021.07.06.451350

**Authors:** Judith C. Birkhoff, Anne L. Korporaal, Rutger W.W. Brouwer, Karol Nowosad, Claudia Milazzo, Lidia Mouratidou, Mirjam C.G.N. van den Hout, Wilfred F.J. van IJcken, Danny Huylebroeck, Andrea Conidi

## Abstract

Perturbation and mechanistic studies have shown that the DNA-binding transcription factor Zeb2 controls cell fate decision and differentiation and/or maturation in multiple cell lineages in embryos and after birth. In cultured embryonic stem cells (ESCs) *Zeb2*’s strong upregulation is necessary for the exit from primed pluripotency and for entering general and neural differentiation. We edited mouse ESCs to produce epitope-tagged Zeb2 from one of its two endogenous alleles. Using ChIP-sequencing, we mapped 2,432 DNA-binding sites of Zeb2 in ESC-derived neuroprogenitor cells (NPCs). A new, major site maps promoter-proximal to *Zeb2* itself, and its homozygous removal demonstrates that *Zeb2* autoregulation is necessary to elicit proper Zeb2-dependent effects in NPC differentiation. We then cross-referenced all Zeb2 DNA-binding sites with transcriptome data from Zeb2 perturbations in ESCs, ventral forebrain in mouse embryos, and adult neurogenesis from the mouse forebrain V-SVZ. While the characteristics of these neurodevelopmental systems differ, we still find interesting overlaps. This contributes to explaining neurodevelopmental disorders caused by *ZEB2* deficiency, including Mowat-Wilson Syndrome.

## Introduction

Zeb2 (also named Sip1/Zfhx1b) and Zeb1 (δEF1/Zfhx1a), the two members of the small family of Zeb transcription factors (TFs) in vertebrates, bind to two separated CACCT (often also to CACCTG E2-boxes) and sometimes CACANNT(G) sequences on DNA via two (between Zeb1 and Zeb2) highly conserved, separated clusters of zinc fingers (1–4). Mutations in *ZEB2* cause Mowat-Wilson Syndrome (MOWS, OMIM#235730) (5–7), a rare congenital disease displaying intellectual disability, epilepsy/seizures, typical facial dimorphism, and often Hirschsprung disease, and multiple other defects in the patients (8–10). Typical are also the delay in developmental milestones and motoric development, and eye and tooth anomalies. Other features are specific craniofacial malformation, sensorineural deafness, and HSCR, which originate from defects in the ZEB2-positive (+) cells of the embryonic neural crest cell lineage. Mutant *ZEB2* alleles have meanwhile been determined for about 350 patients (11–16). Reports have described malformations in the central nervous system (CNS) of MOWS patients over broad age range, in which the observed defects locate to the corpus callosum and/or hippocampus, and can be seen by neuroimaging and follow-up of electro-clinical defects, which include focal seizures (16–18).

Zeb2’s action mechanisms, partner proteins, few proven or candidate direct target genes, and genes whose normal expression depends on intact Zeb2 levels, have been studied in various cell types, explaining specific phenotypes caused by Zeb2 perturbation. Zeb2 DNA-binding around candidate direct target genes helped to explain *Zeb2* loss-of-function phenotypes in ESCs and cells of early and late embryos, and in postnatal and adult mice. These genes are involved in ESC pluripotency (*Nanog, Sox2*), cell differentiation (*Id1*, *Smad7*), and maturation of various cell types, e.g., in embryonic cortical and adult neurogenesis (*Ntf3, Sox6*), as well as epithelial-to-mesenchymal transition (EMT) (*Cdh1*) (19–26). In reverse, subtle mutagenesis of Zeb2 DNA-binding sites in demonstrated target genes has confirmed Zeb DNA-binding and its repressive activity (e.g., on mesodermal X*Bra* (27) and epithelial *Cdh1* (19)).

Despite its critical function in the precise spatial-temporal regulation of expression of system/process-specific relevant genes during embryogenesis and postnatal development, but recently also adult tissue homeostasis and stem cell-based repair, and acute and chronic disease (28, 29; for a recent review, see (30)), ChIP-sequencing (ChIP-seq) data for Zeb2 has been obtained in very few cases only. A major reason is that ChIP-seq grade antibodies specific for Zeb2 are not readily available. Data have been published for high-Zeb2 hepatocellular carcinoma and leukemia cell lines, or cultured cells that overproduce tag-Zeb2 from episomal vectors or the safe *Rosa26* locus (25, 31, 32), neither of which represent endogenous Zeb2 levels nor express *Zeb2* with normal dynamics. Furthermore, most, if not all, anti-Zeb2 antibodies cross-react with Zeb1, so do not discriminate between both proteins when their presence overlaps or succeeds to one another. However, they compete for the same target genes, which for the individual proteins depends on cell identity/state, extrinsic stimulation or cellular context (e.g., in somitogenesis (33) and melanoma (34)).

During neural differentiation (ND) of mouse (m) ESCs, Zeb2 mRNA/protein is undetectable in undifferentiated cells, whereas its strong upregulation accompanies efficient conversion of naïve ESCs into epiblast stem cell like cells (EpiLSCs) and is essential for subsequent exit from primed cells and onset of differentiation, including progression to neuroprogenitor cells (NPCs) (25). We have edited one *Zeb2* allele of mESCs by inserting a Flag-V5 epitope tag (in brief, V5) just before the stop codon, in-frame with the last exon (ex9 of mouse *Zeb2*, (35)). Such Zeb2-V5 mESCs were then differentiated to NPCs, and Zeb2 binding sites were determined by V5-tag ChIP-seq. Doing so, we identified 2,432 binding sites, 2,294 of which map to 1,952 protein-encoding genes. We then cross-referenced the ChIP+ protein-encoding target genes with RNA-seq data of deregulated expressed genes (DEGs) in cell-type specific neurodevelopment-relevant Zeb2 perturbations (23, 36, 37). Although we compare non-identical systems, the overall approach still revealed a number of interesting overlaps, as well as Zeb2’s role in regulating critical targets in neurodevelopment. Taken together, we report for the first time the identification of endogenous genome-wide binding sites (GWBS) in ESC-derived neural cells for the MOWS TF ZEB2.

## Results

### Heterozygous Zeb2-V5 ESCs differentiate as wild-type cells

Addition of short epitope(s) at the N- or C-terminus, as well as activation/repression domains of heterologous TFs at the Zeb2 C-terminus, does not interfere with Zeb2’s DNA-binding (as tested in *Xenopus* embryos (38), heterologous cells (39), mouse forebrain (36) and mESCs (25)). Here, we have used a CRISPR/Cas9 approach (see Materials & Methods) to insert an in-frame Flag-V5-tag encoding sequence in *Zeb2*-ex9 of mESCs (clone 2BE3; Fig. S1). Allele-specific RT-qPCR using primers that amplify sequences between the ex9 and the V5-tag showed mRNA expression from the tagged allele during ND in cell culture at day (D) 0, 4, 6 and 8 (Fig. 1A). Western blot analysis in nuclear extracts of ND-ESCs at D8, thus in NPCs (25), confirmed the presence of Zeb2 of expected relative molecular mass, using either anti-V5 (αV5) or anti-Zeb2 antibodies (Fig. 1B). Both the Zeb2-V5 and wild-type ESCs were then also verified during ND differentiation for temporal expression of *Zeb2*, core pluripotency genes (*Pou5f1*, *Nanog*, both downregulated upon ND, and *Sox2*, also a NPC TF) and an acknowledged NPC marker (*Pax6)* (Fig. 1C,D).

**Figure 1.**
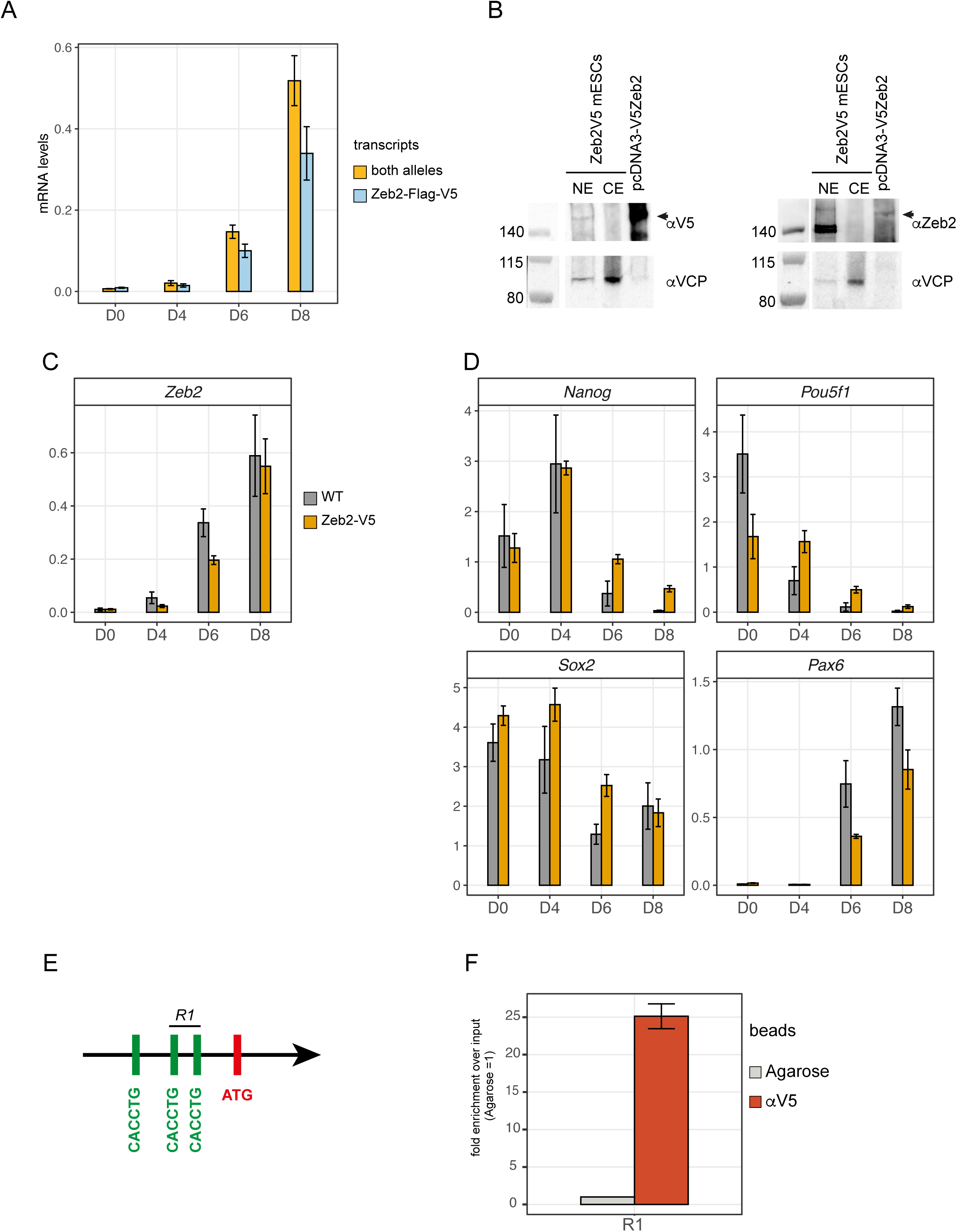
Characterization of the heterozygous Zeb2-Flag-V5 mESC line and ChIP-qPCR validation. A) Allele-specific RT-qPCR using two sets of primers located either in exon7 (and therefore able to detect the whole Zeb2 mRNA produced by both alleles; orange bar) or located in exon9 and the V5-tag (thus recognizing specifically the knocked-in tagged allele; light blue bar). B) Western blot analysis showing V5 epitope containing Zeb2 in ES cell derived NPCs (at D8 of neural differentiation, ND) in nuclear extracts (NE), but not in cytoplasmic extracts (CE). Membranes were blotted with anti-V5 antibody (left panel, αV5) or anti-Zeb2 antibody (right panel, αZeb2 (20). As control, a fraction of Zeb2-rich extract obtained from HeLa cells transfected with pcDNA3-V5Zeb2 vector was also separated in the same gel. C) Zeb2 mRNA levels in wild-type (WT, orange bar) and Zeb2-V5 (grey bar, clone 2BE3, indicated as Zeb2-V5) mESCs during ND. D) Pluripotency markers *Nanog*, *Pou5f1* (*Oct4*) and *Sox2* are down regulated in Zeb2-V5 mESCs similarly to WT. The neuronal marker *Pax6* is also significantly upregulated during differentiation, like in WT mESCs. E) Scheme of the mouse *Cdh1* promoter showing the three E-boxes located upstream the ATG start codon. Zeb2 binds specifically to only two of these, indicated as R1 (25). F) ChIP-qPCR showing enrichment for Zeb2-V5 binding to the R1 region of the *Cdh1* promoter. Agarose beads were used as negative control (in grey).

Both cell lines displayed comparable expression dynamics of these genes, indicating that Zeb2-V5 NPCs at D8 of ND can be used for ChIP-seq. Further confirmation came from selective pull-down of Zeb2 on the known target *Cdh1*, using ChIP-qPCR. Zeb2 binds to 2 of 3 E-boxes in the mouse *Cdh1* promoter (Fig. 1E), which it represses during EMT (19, 25). A ±25-fold enrichment for Zeb2-V5 was obtained when probing this *Cdh1* region using anti-V5 antibody (αV5) conjugated beads compared to agarose beads as negative control (Fig. 1F). Hence, the Zeb2-V5 protein binds to known Zeb2 target sites, and the NPCs are suitable for endogenous mapping of the Zeb2 GWBS.

### One-third of 2,432 Zeb2 DNA-binding sites map close to the transcription start site of transcriptome-confirmed, system-relevant protein-encoding genes, including *Zeb2* itself

αV5-precipitated samples from upscaled Zeb2-V5 NPCs were used for cross-link ChIP-seq, followed by analysis with Galaxy Software (see Materials and Methods; 40). Of the total of 2,432 significant peaks, 2,294 peaks (94% of total) mapped to 1,952 loci that encode protein, while 125 peaks (5% of total) map to micro-RNA (miRNA) genes, and 1% to regions lacking annotation (NA, using ENSEMBL-GRCm38.99; Fig. 2A; File S1). In addition, about 37.5% of all binding sites of Zeb2-V5 are located within −10/+10 kb of annotated transcription start sites (TSS, Fig. 2B). Gene ontology (GO) pathway enrichment analysis of the aforementioned 1,952 loci revealed binding of Zeb2 to classes of genes annotated to Wnt, integrin, chemokine/cytokine (predominantly as defined in inflammation) and cadherin signaling, respectively, as well as to developmental signaling by EGF, VEGF, TGFβ and FGF family pathways (Fig. 2C). Among these 1,952 loci, those for genes encoding transcription regulatory proteins, and post-translational modification as well as metabolic enzymes are well-represented (Fig. 2D).

**Figure 2.**
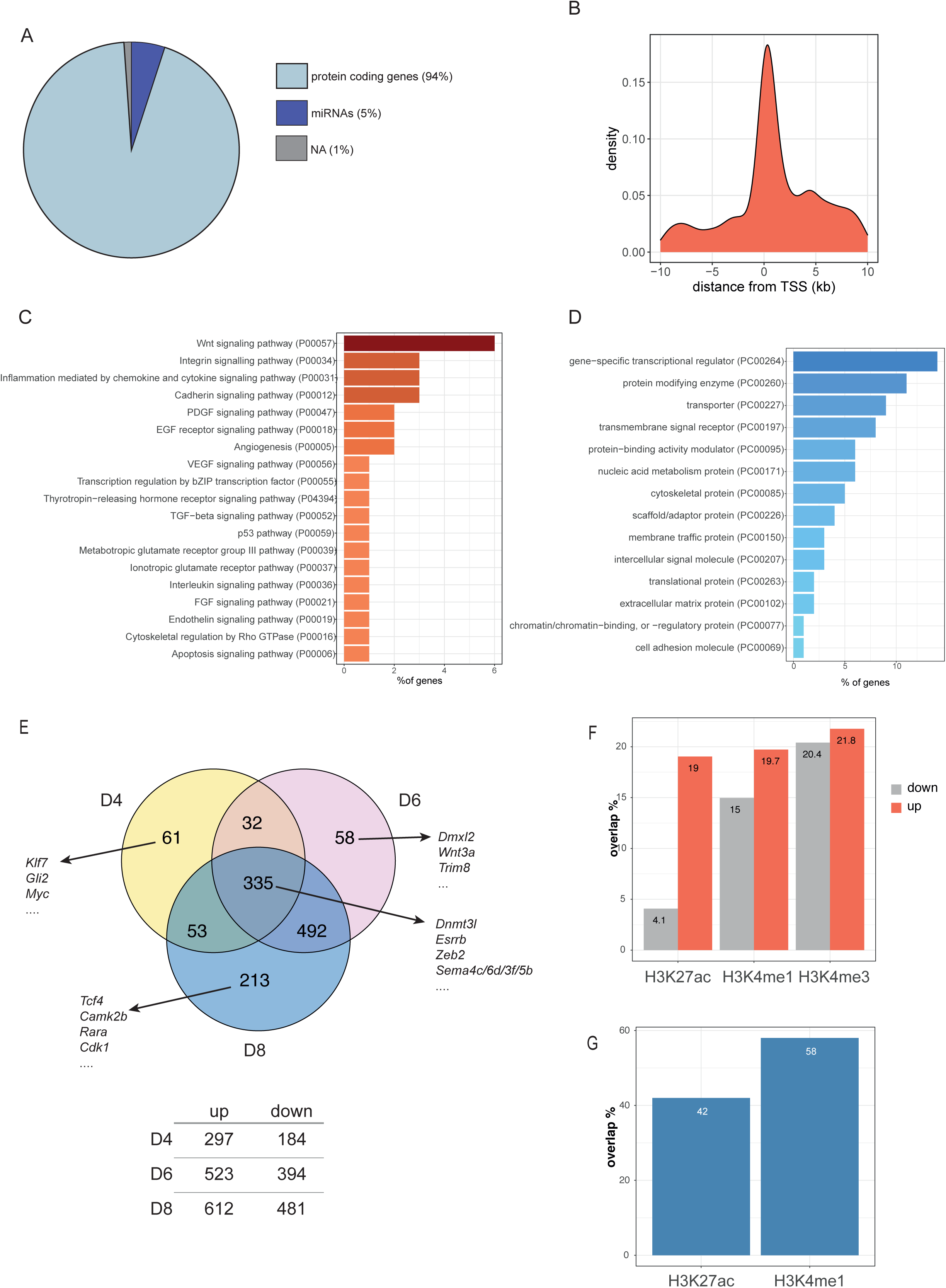
Zeb2V5 protein is mainly recruited at the TSS of transcriptional regulator encoding genes, predominantly those classified in Wnt signaling. A) 2,432 peaks were selected from our ChIP-seq data set (see Materials and Methods). Of these, 94% are associated with protein coding loci, 5% with miRNAs and the remaining 1% map to regions without functional annotation (NA). B) Frequency plot showing the binding of Zeb2V5 at and around (−10 to +10 kb) the TSS. C) The 2,294 peaks map to 1,952 protein encoding genes, many of which operate in Wnt signaling (File S1) or are (shown in D) transcriptional regulators. E) Of the 1,952 protein encoding genes, 1,244 are differentially expressed during ND, when compared to the undifferentiated state (at D0). Of these 1,244 genes, 335 are differentially expressed at all three time points of ND. A few examples are listed of DEGs uniquely expressed at one time point, as well as these that are shared among three time points; a full list is provided in File S2. F) Overlap of the Zeb2 bound regions with H3K27ac, H3K4me1 and H3K4me3 histone marks in the −10/+10 kb from the TSS of up- or down-regulated genes at D8 of mESCs differentiation. G) Overlap of the Zeb2 bound regions outside the −10/+10kb region from the TSS with histone marks.

In parallel we applied bulk temporal RNA-seq of wild-type mESCs at D0 (undifferentiated), D4 (neural induction), D6 (early NPCs) and D8 (NPCs) and checked the expression dynamics of the 1,952 Zeb2-bound genes (from the D8 sample). Among these, 1,244 were subject to significant transcriptional regulation between D4-8 as compared to D0 (Fig. 2E; log2FoldChange <-0.5 or >0.5 and p-value < 0.05; low-stringency analysis was opted to assess also small differences in mRNA of Zeb2-bound genes). Further, 335 of these genes are commonly expressed between D4-6-8, but at different levels, including *Zeb2* itself (for lists of all DEGs, see File S2). Fig. S2A depicts the D4, D6 and D8 transcriptomes of ND-mESCs, each compared to D0, with indication whether the genes are bound or not by Zeb2, as determined by Zeb2-V5 ChIP-seq. At each of these respective time points, hence at different Zeb2 mRNA level, about 12% of the DEGs are bound by Zeb2 (Fig. S2B).

Among the Zeb2-bound genes that normally become down-regulated, *Dnmt3l* and *Esrrb* are present, suggesting that upregulation of *Zeb2* in ND-ESCs (D6 and D8) directly causes downregulation of these two genes accordingly (Fig. S2C; File S2). Zeb2 has been suggested as direct repressor of *Dnmt3l* and *Esrrb*, facilitating the switch from self-renewal of ESCs to their exit from pluripotency, and promoting differentiation. Indeed, expression levels of all *Dnmt3* genes remained higher in *Zeb2*-knockout (KO) ND-ESCs (25). However, these *Zeb2*-KO cells also convert very inefficiently into EpiLSCs and fail to exit from primed pluripotency. Importantly, among the Zeb2-binding genes whose mRNA levels increased during ND, *Zeb2* itself is also present (yellow dot, Fig. S2C), indicating autoregulation. In fact, in this ND model the highest recruitment of Zeb2-V5 in ChIP-seq data was mapped upstream the TSS of *Zeb2* (File S1).

Out of the 1,244 Zeb2-bound genes that significantly changed steady-state mRNA levels in our D4 to D8 transcriptome data sets, 213 are exclusive DEGs in NPCs at D8 (Fig. 2E; Fig. S2D). Among these, *Tcf4* is bound by Zeb2 and becomes highly upregulated in NPCs (Fig. 2E; Fig. S2D). Tcf4 is a ubiquitous basic helix-loop-helix (bHLH) type TF that binds to E-boxes, and co-operates in many isoforms (41, 42) with cell-type specific bHLH TFs as heterodimers, which are active during CNS development ((43, 44); for a review, see (45)). In oligodendrocyte precursors (OPCs), Tcf4 is essential for their subsequent differentiation. It dimerizes with the lineage-specific bHLH-TF Olig2, further promoting their differentiation and maturation (46), while Zeb2 together with upstream Olig1/2 are essential for myelinogenesis in the embryonic CNS (21). Here, Zeb2 generates anti-BMP(-Smad)/anti-Wnt(-β-catenin) activities, which is crucial for embryonic CNS myelinogenesis emanating from differentiation of OPCs. The regulatory action of Zeb2 on the *Tcf4* target gene, as found in mouse cells by our ChIP-seq, may underpin phenotypic similarities between MOWS and Pitt-Hopkins syndrome patients (PTHS, OMIM #610954), the latter caused by mutations in *TCF4* (47), making us speculate that *TCF4* may be deregulated in MOWS neural cells.

### Zeb2 peaks overlap with active enhancers and promoters within −10/+10 kb from the TSS

To assess whether Zeb2-peaks are present in the regulatory regions of up- or down-regulated genes from our transcriptome data, we cross-referenced the coordinates of the Zeb2 broad peaks within − 10/+10kb from the TSS, with the mouse ChIP-seq datasets available in ENCODE for neuronal tissues (cerebellum, cortical plate, olfactory bulb, forebrain, midbrain, hindbrain, neural tubeolfactory bulb). We found that of the different histones ChIP-seq datasets available in ENCODE H3K27ac, H3K4me1 and H3K4me3 marks were overlapping with our ChIP-seq data (Figure 2F). The H3K27ac signature strongly overlaps with the Zeb2 peaks in genes upregulated at D8 (19% in upregulated genes vs. 4% in downregulated genes). For H3K4me1 and H4K4me3 marks, no big difference in overlap between up- and down-regulated genes was observed. While H3K27ac mark is associated with active enhancers, H3K4me1 is associated with primed enhancers, and H3K4me3 is considered a “promoter” marker (48). Taken together these data suggest an activating role for Zeb2 at this stage of differentiation. Outside the −10/+10kb considered range, about 48% of the identified peaks overlap for about 48% with H3K27Ac and 52% with H3K4me1 histone marks (Fig. 2G).

We then performed motif enrichment analysis using UniBind (https://unibind.uio.no) for TFs that could bind the Zeb2-bound peaks and in proximity of up- or down-regulated genes at D8 of mESCs differentiation (S3A and B). In those peaks close to the TSS of upregulated genes, motifs for Sox2, Gata2 and Tcf3 are very abundant. These TFs are known to have a function during NPC or neural differentiation. Sox2 is an acknowledged marker for neurogenesis (49) and during later stages of differentiation. It has been demonstrated that Zeb2 exhibit anti-Sox2 activities in myelination and re-myelination by adult Schwann cells in the PNS, needed for normal progression of commitment, differentiation and maturation in this glial cell lineage (21, 24, 29,50). Tcf3 (also named E2A) plays a role in stem cell self-renewal (51), but is also important during neural fate commitment and possibly repressing Nodal signaling during neural differentiation (52). Gata2 has been associated with negatively regulating proliferation in NPCs and, by doing so, directing these cells further into differentiation (53). However, how Zeb2 acts upon, or together with these TFs during NPC differentiation. is not fully worked-out yet.

In the peaks close to the TSS of downregulated genes there is a prevalence for CTCF, Fos, Myc and Stat5a binding. CTCF is likely of specific interest for its moieties to act as a link between 3D genome architecture to gene regulation. During NPC differentiation however, it was observed that 40% of the NPC-specific DNA-loops were not CTCF-dependent, whereas in other cell-state specific loops, this was only 10%, indicating a less important role for CTCF in the regulation of NPC differentiation compared to differentiation into other cell lineages (54). This might indicate an interesting role for Zeb2 in repressing CTCF levels during NPC differentiation, supporting the subsequent activation of NPC-specific genes. Also here, more studies are required to get more insights into the co-operativity or counteracting actions of Zeb2 with candidate TFs in the regulation of the candidate Zeb2 targets.

### Meta-analysis of RNA-seq data from selected neural-system *Zeb2* perturbations and the NPC ChIP-seq data reveal interesting overlaps

We performed a meta-analysis of three published transcriptome data sets from control and *Zeb2*-KO mice: sorted E14.5 mouse ventral forebrain interneurons (Nkx2.1-Cre driven *Zeb2*-KO; (36)) and sorted (at P2) progenitors of the V-SVZ, an adult neurogenic niche (Gsh2-Cre; 36). In addition, we used high-throughput RT-qPCR data generated on a Fluidigm platform and obtained after esiRNA-based knockdown (KD) of Zeb2, as part of a systems-biology study in ND-mESCs ((37) (the Zeb2 KD data subset was kindly provided by R. Dries). From these respective datasets, the DEGs upon the Zeb2 perturbations (p-value <0.05; log2FoldChange<l-1 and >1) were filtered. This identified (i) genes that depend on normal Zeb2 levels for their downregulation/repression (if directly by Zeb2, as repressor) or (ii) other genes that depend on Zeb2 for their upregulation/activation (if directly by Zeb2, as activator) (for Zeb2 as dual TF, see (29); (30, 55)). In parallel, our 2,294 Zeb2-V5 sites mapping to the 1,952 protein-encoding genes were filtered from the complete ChIP-seq dataset, and then used as reference for the RNA data sets (Fig. 3A).

**Figure 3.**
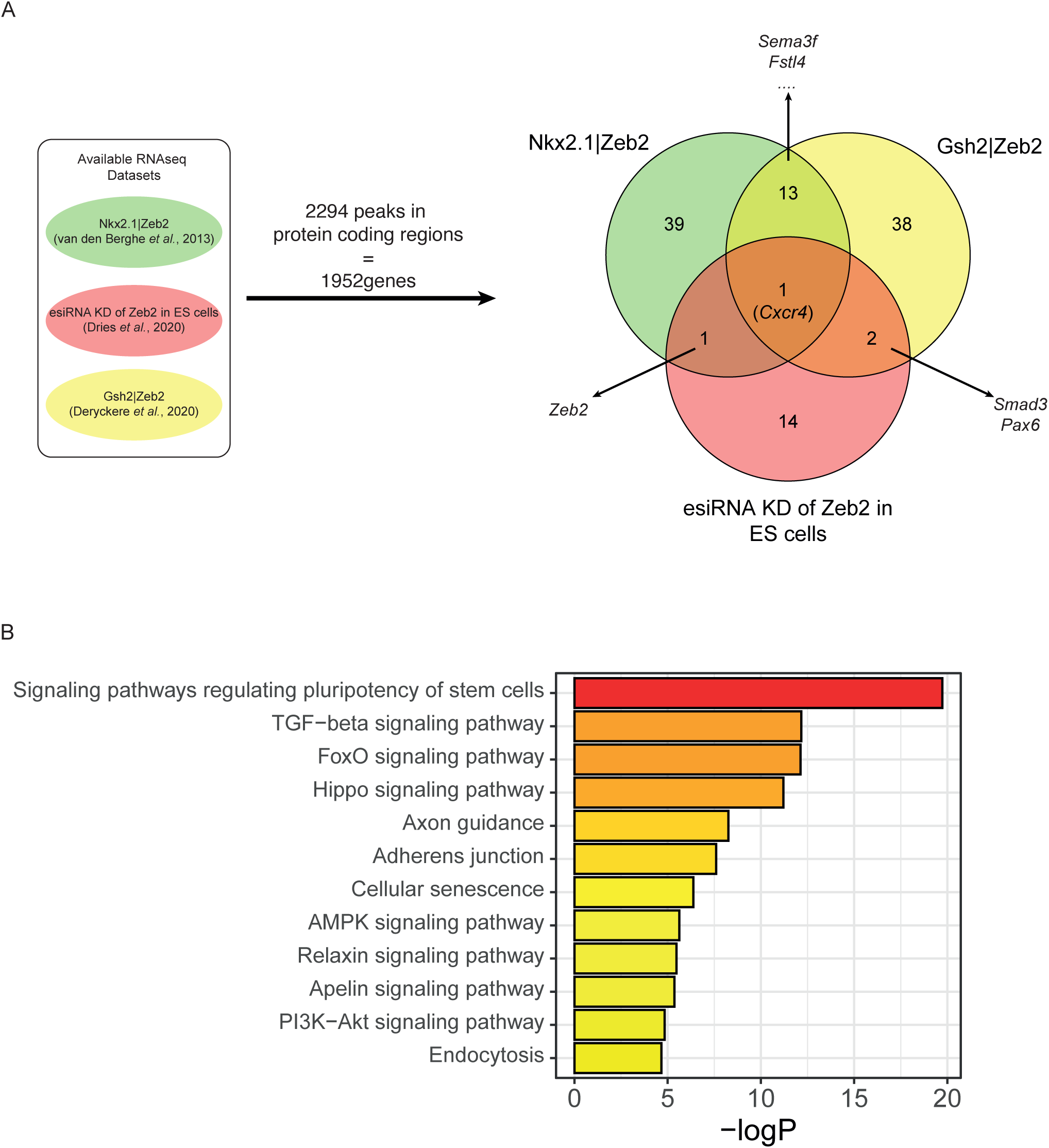
Schematic representation of the meta-analysis Zeb2-bound genes versus RNA-seq datasets. A) 108 genes bound by Zeb2 are also differentially expressed in the three datasets from other studies in mouse models and ESCs. *Cxcr4* is the only DEG bound by Zeb2 and common among the three datasets. B) These 108 genes mainly map to signaling pathways regulating stem cell pluripotency, and effects of TGFβ family, FoxO, and Hippo signaling/activity.

This cross-referencing identified 108 protein-encoding genes among the three transcriptomic data sets and ChIP-seq data set (Fig. 3A). Fig. S4 shows a heatmap of the changes in mRNA levels of these 108 genes during ND of wild-type ESCs and their correlation with the analyzed datasets. Noteworthy, only *Cxcr4* was common to all RNA data sets. This is likely due to the fact that two RNA-seq sets are generated in different brain/neuron cell-type *in vivo* mouse models, while the other steady-state RNA level data documented the effects of Zeb2-KD on mRNA levels of (only 96 in total) TGFβ/BMP-system components (37), so the timing does not completely overlap with our ChIP-seq dataset. However, Cxcr4 and its ligand Cxcl12/Sdf-1 are crucial for migration of interneurons from the ventral forebrain to the neocortex (56, 57), processes co-controlled by Zeb2 as shown in cell-type specific KO mice (36). Furthermore, the identified 108 genes are involved in regulation of stem cell pluripotency, signaling by TGFβ, FoxO, and Hippo, and in axon guidance. Taken together these data further confirms the pivotal role of Zeb2 in these processes.

### Zeb2 directly controls TGF**β**/BMP-system component and neuronal differentiation/ migration genes

We then validated 14 out of the 108 cross-referenced target genes, selected based on either being already known as a target of Zeb2 (*Nanog*), or as TGFβ/BMP-system component (*Bmp7*, *Tgfbr2*, *Smad1*, *Smad2*, *Smad3*, *Id2*, *Cited2*), or having a crucial role in neurogenesis and neuronal maturation (*Sema3f, Cxcr4*, *Lhx5*, *Ntng2*, *Pax6*, *Tcf4*; their mRNA levels in ND-ESCs are highlighted in the heatmap in Fig. S4). Because *Zeb2*-KO ESCs do not exit from primed pluripotency and thus cannot differentiate (25), we chose to validate our findings using shRNA-mediated Zeb2-KD at ND-D8, and analyzed these aforementioned 14 genes two days later (at D10 NPCs) (Fig. 4A); at this read-out time point >50% reduction of Zeb2 mRNA was obtained (Fig. 4B).

**Figure 4.**
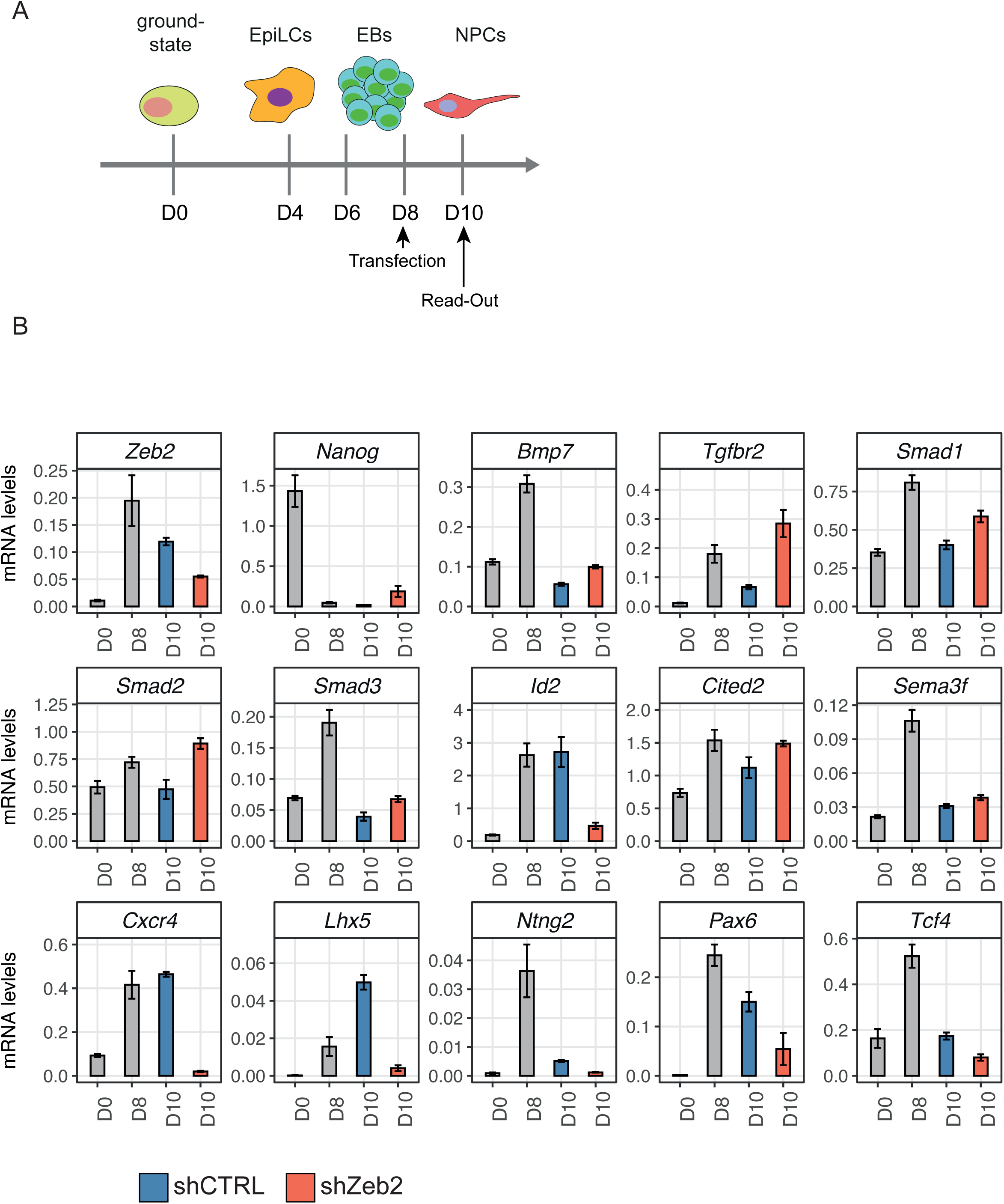
shRNA-mediated KD of Zeb2 discriminate between primary and secondary target genes. A) Schematic overview of the shRNA transfection targeting Zeb2 (shZeb2) and read-out of the effect. Cellular aggregates at D8 of ND are dissociated and transfected with shZeb2 or against a scrambled, control sequence (shCTRL). Read-out is done two days after the start of shRNA addition. The list of shRNAs is given in Table S2. B) *Zeb2* levels after KD were reduced to 40-50% of their normal level (shZeb2, orange bars) compared to shCTRL (blue bars). *Bmp7*, *Cited2*, *Nanog*, *Sema3f*, *Smad1*, *Smad2*, *Smad3*, and *Tgfbr2* were significantly upregulated following Zeb2 KD, whereas genes encoding for neuronal specification and migration (*Cxcr4*, *Lhx5*, *Ntng2*, *Pax6* and *Tcf4*) were downregulated.

Zeb2-KD resulted in reduced mRNA levels of *Cxcr4*, *Ntng2* and *Pax6* (Fig. 4B), genes that are each involved in neuron specification and migration (58, 59). Zeb2-KD also caused down-regulation of *Lhx5,* involved in differentiation of interneurons, including cytoskeletal rearrangements during dendritogenesis (60), and of *Tcf4*, which acts in neurogenesis (43, 44). *Sema3f is a* cue for axon outgrowth and neuron migration guidance, and was slightly upregulated (Fig. 4B). These results confirm the regulation by Zeb2 of its direct targets in later phases of neuronal differentiation/migration.

The expression of *Nanog*, the promoter of which binds Zeb2 as a repressor (25), was increased in the Zeb2-KD cells (Fig. 4B). Zeb2-KD caused increased mRNA of *Bmp7*, *Tgfbr2, Smad1*, *Smad2, Smad3* and *Cited2* (Fig. 4B), fitting with the normal levels of Zeb2 that mount anti-TGFβ/BMP family effects (29). In contrast, *Id2* is significantly downregulated in shZeb2-treated ESCs (Fig. 4B). *Id2* is normally activated by BMP-Smads and, together with other Id proteins (Id1, Id3 and Id4), inhibits cell differentiation, e.g., Zeb2 represses *Id2* in immune cells to promote differentiation (22). However, *Id2* as well as other *Id* genes (61–63) is, like *Zeb2* (20, 36), also expressed in the developing forebrain. Taken together, these data suggest an active and direct role for Zeb2 in repressing genes regulating stem cell pluripotency as well as a number of TGFβ/BMP-system components (*Bmp7*, *Smad1/2/3*), but also in activating genes during neurogenesis (*Cxcr4, Ntng2, Lhx5*).

### Zeb2 potentiates its own gene expression, which is crucial for proper control of some of its direct target genes

Strikingly, in our ChIP-seq dataset, the peak with the highest enrichment (∼200-fold) mapped 232 bp upstream of the Zeb2 TSS (Fig. 5A, File S1). For further functional studies of this site, we deleted the encompassing region (chr2:45109746-45110421) using CRISPR/Cas9 in wild-type mESCs, thereby obtaining *Zeb2^ΔP/ΔP^* ESCs (Fig. S6; see Materials and Methods). Fig. 5B shows that the Zeb2 mRNA levels in the homozygous *ΔP* clone stayed significantly low during ND, already from D8 onwards, compared to control cells, classifying this *Zeb2^ΔP/ΔP^* clone as an alternative Zeb2-KD.

**Figure 5.**
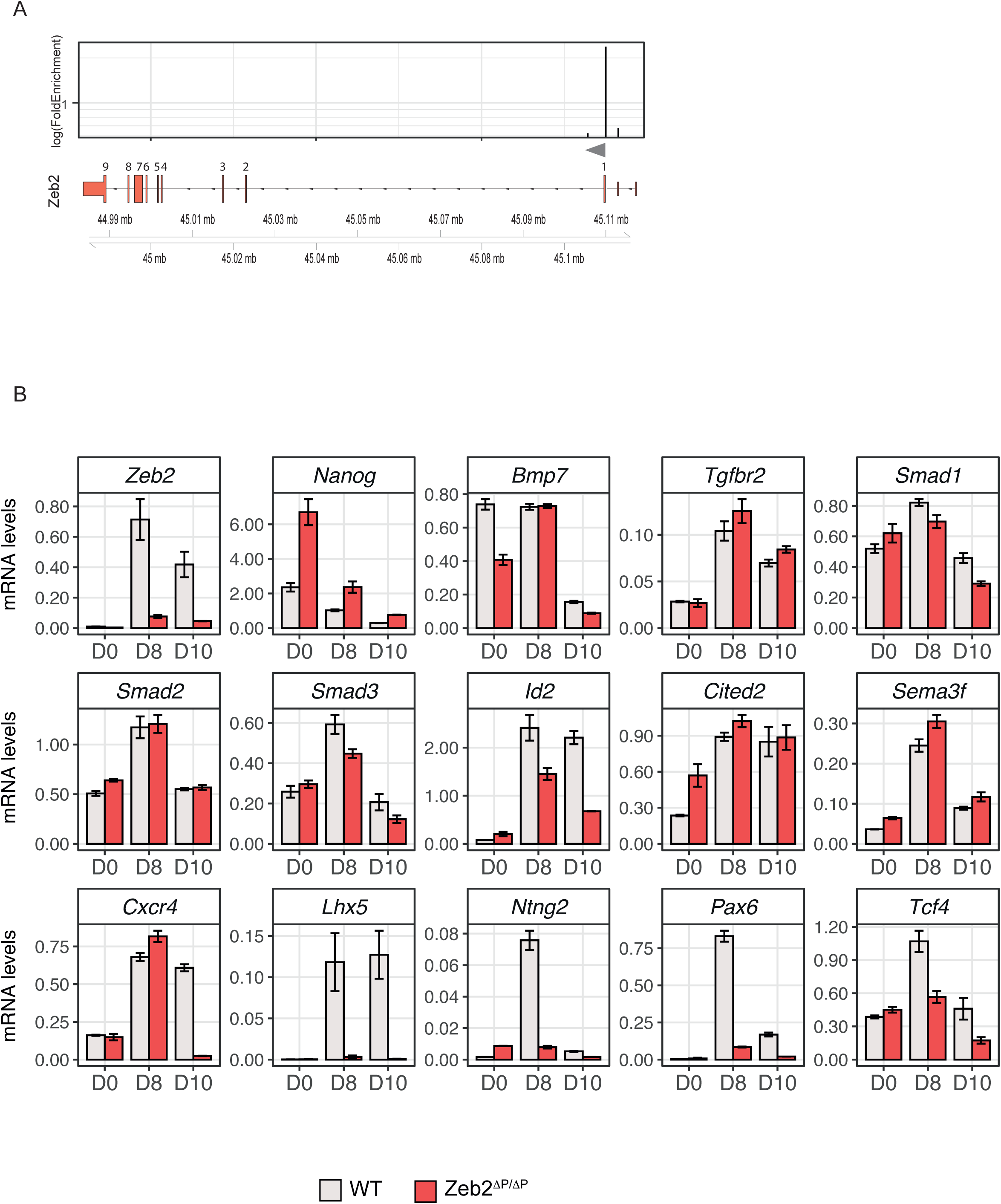
Deletion of the Zeb2-binding, candidate autoregulatory site (chr2:45109746-45110421) impairs Zeb2 mRNA levels and neuronal markers. A) Schematic overview of the log(FoldEnrichment) of the peaks identified by ChIP-seq located in the mouse *Zeb2* locus, and localization on top of *Zeb2* intron/exon structure. The highest peak is located 232 bp upstream of the first translated exon. Grey arrow indicates the TSS of *Zeb2*. B) Zeb2 mRNA levels are strongly reduced in the *Zeb2^ΔP/ΔP^* clone. Expression levels (mRNA) of the target genes validated with shRNA (Figure 4). Most, but not all of the genes found to be affected following Zeb2 KD are also deregulated in *Zeb2^ΔP/ΔP^* mESCs in particular *Id2* and the neuronal markers *Cxcr4*, *Lhx5*, *Ntng2*, *Pax6* and *Tcf4*.

We then used these *Zeb2^ΔP/ΔP^*mESCs to read-out the same genes that depend on intact Zeb2 levels and are Zeb2 ChIP*+* (see Fig. 4B). Levels of Zeb2 mRNA stayed abnormally very low at D10 in *Zeb2^ΔP/ΔP^* ND-mESCs, whereas *Nanog* was still expressed and remained higher than in control WT cells (Fig. 5B). Hence, Zeb2 levels, including those achieved by autoregulation, are critical, but to a different degree for sets of genes. The latter include neuronal-relevant genes such as *Cxcr4*, *Lhx5*, *Ntng2*, *Pax6*, *Tcf4* (Fig. 5B). Among the TGFβ/BMP-system components (see Fig. 4) we observed a slight reduction of *Bmp7*, *Smad1* and *Smad3*, whereas *Tgfbr2*, *Smad2*, *Cited2* and *Sema3f* expression was not affected in *Zeb2^ΔP/ΔP^* ND-mESCs. We speculate that Zeb2, and now including its newly identified autoregulation, plays an active role in regulating a number of genes involved in neuron determination and, together with other TFs, TGFβ/BMP-system component genes. Thus, the precise amounts of Zeb2, and in a critical stage also its autoregulation, are crucial in discriminating genes where Zeb2 plays the aforementioned primary, active role (as for *Cxcr4*, *Lhx5*, *Ntng*2 etc.). For these genes, ∼50% reduction or the novel Δ*P/*Δ*P* “peak” mutation is sufficient to strongly deregulate them, but other genes’ expression is either not or just slightly affected (*Sema3f*, *Smad2*, *Cited2* vs. *Bmp7*, *Tgfbr2*, *Smad1*, *Smad3*).

Because Zeb2 also binds phospho(p)-Smads (4, 21, 23, 29), we also scanned the Zeb2 ChIP+ direct target genes, and for which we saw strong deregulation upon Zeb2-KD and/or in *Zeb2^ΔP/ΔP^* cells during ND (wherein notably Smad-activation is not stimulated), for the presence of (i) the Zeb half-sites CACCT(G) (pragmatically neglecting the variable spacing between them; 64) that combine with (ii) candidate p-Smad binding and responsive genes (using GTC(^T^/_G_)CT(^T^/_G_)(^A^/_C_)GCC for p-Smad1/Smad5, GTCTAGAC for p-Smad2/3) and (iii) the co-Smad Smad4 (C(^C^/_T_)AGAC) (65), using the Jaspar database. Fig. S6 shows the distribution of such identified Zeb and Smad-binding motifs (threshold score >85%) in those genes strongly affected by Zeb2-KD and/or in *Zeb2^ΔP/ΔP^* cells. Interestingly, in the regions where Zeb2 binds close to the TSS (*Zeb2*, *Ntng2*, *Lhx5, Nanog*), the p-Smad and Smad4 binding elements are sometimes present in very close proximity of the ChIP*+* Zeb2- bound E-box, indicating a possible cross-talk between receptor-activated Smads and Zeb2 in regulating target genes.

## Discussion

We report for the first time the endogenous GWBS for Zeb2 by ChIP-seq, in ESC-derived NPCs gene-edited for this. We have previously used ESCs established from *Zeb2*^Δ^*^ex7/^*^Δ^*^ex7^*-KO (66) pre-implantation embryos and, for rescue purposes, such KO ESCs cells in which Flag_3_-Strep-Zeb2 was produced from (a Cre-controllable) *Rosa26* locus (25). Conceptually, with regard to Zeb2 levels, the latter cells are different from the mESCs that were newly established here, since in the original Rosa26-Zeb2 cells *Zeb2* is not subjected to its normal temporal regulation during cell differentiation.

However, precise dosage of Zeb2 is a critical factor *in vivo* (for a recent discussion, see 30). This is concluded from transgenic Zeb2 cDNA-based rescues in *Zeb2*-KO ESCs and similar genetic rescues in *Zeb2*-mutant cells in mice, which can via heterozygous/homozygous combinations create an elegant and large panel of Zeb2 levels (in interneurons (36), NK cells (67), ESCs (25)). Another illustration of the relevance of fine-tuned control of Zeb2 levels are miRs targeting Zeb2, and lncRNAs that regulate these miRs (68–71), with Zeb2 on its turn also controlling some of its own miR-encoding genes or clusters (72–74). In our ChIP-seq we find 125 peaks (∼5% of the total) that correspond to the TSSs of 98 miR-genes (File S1). Among these miR-genes, Zeb2 binds to loci encoding miR-144, miR-148a, miR-9 and miR-153, known to target Zeb2 in the context of e.g., tumor progression (75–78). We recently added the identification, in human iPSCs subjected to ND, of *ZEB2* distant (∼600 kb upstream) enhancers, which act through DNA-looping to the *ZEB2* promoter-proximal region (79). In addition, we have documented dynamic expression patterns of *Zeb2* in early embryos (27, 33, 80–82). We have also shown that cDNA-based expression of various tag-Zeb2 proteins is compatible with functional embryology-type and action mechanism studies (25, 38, 39). Importantly, our Zeb2-V5 allele enables normal production of tag-Zeb2 for the first time from its endogenous locus.

Only two studies present ZEB2 ChIP-seq data in human cells, i.e. SNU398 hepatocellular carcinoma and K562 erythroleukemia cells, respectively (31, 32). In K562 cells ZEB2 binds to the promoters of *NR4A2*, *NEUROG2* and *PITX3*, expressed in midbrain dopaminergic neurons, wherein Zeb2 negatively regulates axon growth and target innervation in mice (83). In SNU398, ZEB2 represses *GALNT3*, which is normally expressed in epithelial cells. This repression coincides with acquisition of a mesenchymal phenotype, linking ZEB2 here again to an EMT-like process. These two valuable studies also present limitations. The use of cancer cell lines of genomic instable nature may create possible bias in ChIP-seq, and in any case they overproduce ZEB2. Our ChIP-seq identifies >2,400 peaks for Zeb2-V5 in mESCs at ND-D8. This is in cells not stimulated with TGFβ family ligands, with 37.5% of Zeb2 sites mapping close to te TSSs (when defined as −10/+10kb). Most of these genes function in growth factor or cytokine signaling and/or encode transcriptional regulators, the latter suggesting that Zeb2 orchestrates other co-operating TFs driving the transcriptomic signature of NPCs. The regulation of Wnt signaling by Zeb2 is in line with observations that inhibition of the Wnt-βcatenin pathway suppresses ND *in vitro* and *in vivo*, and that Wnt (and Zeb2)-controlled *Tcf4* expression promotes neurogenesis and is required for normal brain development (84–88).

The 2,294 Zeb2 peaks map to 1,952 protein-coding genes, of which 1,244 are DEGs in ND-mESCs. Strikingly, the strongest enrichment of Zeb2 occurs on the *Zeb2* promoter itself, leading to the identification of a novel self-regulatory mechanism where Zeb2 binds upstream its TSS to maintain its levels sufficiently high, also and at least during ND. While this autoregulation needs further investigation in Zeb2-dependent differentiation and/or maturation of other cell types (e.g. in cKO mouse models or in ND-iPSCs derived from appropriate MOWS patient cells), we propose that lower Zeb2 levels might compromise this autoregulatory loop. Deletion of the autoregulatory site from both *Zeb2* alleles (in the Δ*P/*Δ*P* cells), results in a significant decrease of Zeb2 mRNA levels but *Zeb2* is still partially expressed, and these cells can still exit from pluripotency and differentiate (contained in part within Fig. 5; *data not shown*). A number of genes, which are mainly linked to neuron maturation, are significantly affected in *Zeb2^ΔP/ΔP^* ESCs, whereas TGFβ/BMP-system component genes are not deregulated. Zeb2 dosage might thus underlie this difference in regulating its direct, ChIP*+* genes in our ESCs. Zeb2 might be key to maintaining expression of neuronal genes, while for TGFβ/BMP-system genes Zeb2 may co-operate with other TFs (including p-Smads) or DNA-modifying enzymes to regulate the expression of target genes.

Zeb2 binds to receptor-activated phospho-Smads (pSmads), and several studies indicate its negative regulation of BMP-Smad activation of specific target genes, although Zeb2 also has Smad-independent functions (23, 29). BMP-pSmads bind to GGCGCC with high affinity (89). Morikawa and co-workers (90) have confirmed these results using ChIP-seq, and identified also a lower-affinity (so, higher BMP-doses required) BMP-Smad element (GGAGCC). For achieving full responsiveness it was proposed that the GG(^A^/_C_)GCC element needs to be coupled with a Smad4 site, ideally located 5 bp away (90). We find that in primary targets affected by varying levels of Zeb2, E-boxes are located close to Smad-binding motifs. However, whether Zeb2 and Smads are co-present in target regions requires further experiments, such as ChiP-on-ChIP assays, and (non-neural) differentiation protocols (involving stimulation of the cells by addition of BMP and/or Nodal). However, these studies may be further complicated because of post-translational modification status of Zeb2, nuclear p-Smads and/or Smad4 (91–94).

Striking are also the 1,093 Zeb2-binding DEGs at D8. When we performed a gene-to-disease association using the human orthologues of these D8-DEGs, we found a clear association with several disorders (Fig. S6). These include neurodevelopmental, mental and ocular defects, which occur in MOWS. Altogether, our data may provide novel insights in MOWS due to suboptimal ZEB2 amounts in patients, and from now in includes an autoregulation aspect, as well as *ZEB2* as a putative modifier gene for many other congenital disorders.

Several *Zeb2*-cKO mouse models have been generated, and for many bulk RNA-seq data are available, from which here we selected two such data sets (23, 36, 95). In addition, similar data were obtained for cultured mESCs, either *Zeb2*-KO cells (25) or cells submitted to ND wherein e.g., Zeb2-KD was performed (37). Unfortunately, we could not include *Zeb2*-KO mESCs in these comparisons, for they convert less efficiently to EpiLSCs and fail to differentiate beyond this EpiLC state (25). The meta-analysis of these three different data sets, overlaid with the 1,952 Zeb2-V5 ChIP+ loci/genes, show therefore a limited number of common targets, *Cxcr4* being the only one common in all data sets. This specifically narrows the Zeb2-bound gene collection to 108 in total. However, and interestingly, the latter enrich for GO terms such as pluripotency of stem cells, signaling by TGFβ and Wnt, cell fate commitment and neuron differentiation, all processes where Zeb2 plays a crucial role.

Out of these 108 genes, we selected 14 covering TGFβ/BMP signaling, pluripotency, neuron migration an differentiation/maturation, and checked their levels 2 days after Zeb2-KD at ND-D8. Most of these 14 genes relevant to NPC status showed to be critically depending on intact levels of Zeb2. They may help to explain why the defects caused by MOWS are observed later after birth and why (the few) missense mutations in MOWS (besides the more abundant significant deletions) present with milder syndromic manifestation. Both the cross-reference of Zeb2-ChIP*+* genes with the transcriptome of ND-ESCs, and the meta-analysis, identify a number of common genes, such as *Bmp7*, *Tgfbr2*, *Tcf4*, *Smad1/2/3*, and *Sema3f* (File S3; Fig. S4), making Zeb2 a likely direct regulator of these genes. It is also intriguing that Zeb2 is recruited to and controlling *Tcf4* at D8, and that *Tcf4* is deregulated upon Zeb2-KD (using esiRNA (37) and shRNA here). Mutations in *TCF4* cause PTHS, a rare neurodevelopmental disorder with some defects overlapping with MOWS. The binding of Zeb2 to *Tcf4* opens new attractive roads to further investigate the crosstalk between these two TFs and their role in regulating crucial aspects of neurodevelopment.

## Materials and Methods

### ESC culture and differentiation

CGR8 (strain 129) wild-type and Zeb2-Flag-V5+ mESCs were cultured and differentiated towards the neural lineage (96, with few modifications). Briefly, mESCs were cultured on 0.1% Gelatin-coated plates in ESC-medium: DMEM supplemented with 15% heat-inactivated (HI) FBS, 2mM L-Glutamine, 1x Non-Essential Amino Acids (NEAA), 143 µM β-Mercapto-Ethanol (β-EtSH) (all ThermoFisher Scientific, TFS) and LIF (10^3^ U/ml).

For ND, 4×10^6^ cells were plated on non-adherent 10-cm dishes (Greiner) and allowed to form cellular aggregates (CAs) in 10 ml CA-medium (DMEM, 10% HI-FBS, 2 mM L-Glutamine, 1x NEAA, and 143 µM β-EtSH). From D4 of ND, cells were grown in CA-medium supplemented with 5 µM Retinoic Acid (RA). During the aggregation stages of ND, medium was changed every other day by carefully collecting the aggregates with a 10-ml pipet and transferring them to a 15-ml conical tube. The CAs were allowed to sink to the bottom of the tube where, after the previous medium was carefully discarded, the CAs were then resuspended in fresh medium and transferred back to the dishes. At D8 of ND, the aggregates were harvested and dissociated by resuspension in 1 ml Accutase (TFS) and pipetting them up-and-down using a 1-ml pipet, after shaking them in a 37 JC water bath for 5 min. The Accutase was deactivated by adding 9 ml of fresh N2-medium (DMEM with 2 mM L-Glutamine, 50 µg BSA/ml and 1x N2-supplement) to the dissociated cells and pelleting the cells gently for 5 min at 200 g. The cells were resuspended in fresh N2. To ensure single-cell suspension, the cells were filtered by passing them through a 40-mm nylon cell strainer (Corning). 2.5 x 10^5^ cells/cm^2^ were plated on Poly-DL-Ornithine hydrobromide (Sigma) / Laminin (Sigma) coated plates. Cells were harvested at D8 or D10 of ND.

### Western blots

To check Zeb2-V5 protein, the 2BE3-clone ESCs were subjected to ND till D8. Cytoplasmic and nuclear fractions were extracted using NePer-kit® (TFS). Protein concentrations were measured using the Bradfort BCA (TFS) and equal quantities of protein lysates were loaded on 6% SDS-PAGs and thereafter cut based on protein relative molecular mass. Gels were then transferred onto nitrocellulose membranes (Amersham Bioscience), which were incubated overnight with anti-Zeb2 (20) and anti-V5 (Life Technologies) antibody, followed by incubation at RT with HorseRadish Peroxidase (HRP) conjugated secondary anti-rabbit and anti-mouse antibodies, respectively (Jackson ImmunoResearch). Protein bands corresponding to Zeb2 or Zeb2-V5 were visualized on an AI-600 digital imager (Amersham|). As loading control, we used Valosin-containing Protein (VCP) and anti-VCP antibody (Santa Cruz sc-57492, mouse).

### RNA extraction and RT-qPCR analysis

Total RNA was extracted from ESCs using TRI Reagent (Sigma), and used for cDNA synthesis with RevertAid RT Kit (TFS) with oligodT-primers. RT-qPCR was performed using SybrGreen dye (BioRad) on a CFX96 T1000 thermal cycler (BioRad). All data shown are averages of 3 independent biological replicates and 3 technical replicates, normalized to β-Actin mRNA levels. Primers are listed in Table 2. Analysis and data visualization was performed in R environment for statistical computing version 3.5.3, implemented with the *tidyverse* version 1.3 package (https://github.com/tidyverse).

### Tag-Zeb2 mouse ESCs

gRNAs (Table 1) targeting *Zeb2*-ex9, and tracrRNA (Integrated DNA Technologies, IDT), were diluted to 125 ng/µl in duplex buffer (IDT). gRNAs were annealed to tracrRNA at 1:1 ratio at 95°C for 5 min and cooling the samples to room temperature (RT, 24°C). 250 ng of these annealed gRNAs were transfected in 350,000 mESCs together with 2 µg pX459-Cas9-puro vector and 1 µg ssDNA oligo of the Donor Template containing the FlagV5-tag sequence (Table 1). Transfection was done in a gelatin-coated 6-well plate using DNA:Lipofectamine2000 (ratio of 1:2). Six hours after transfection the medium was refreshed, at 24 hours the cells were Puromycin-selected (2 µg/ml). After 2 days, the remaining cells were transferred to gelatin-coated 10-cm dishes and given fresh ESC-medium (see below). Per dish 1,000; 1,500; or 2,000 cells were plated and allowed to form colonies. Medium was changed every other day. Colonies were picked, expanded and genotyped by PCR (both outer and inner primer sets were used (Table S1; Fig. S1). All candidate clones were validated by Sanger-sequencing; correct clones were expanded and validated by Western blot.

**Table 1:**
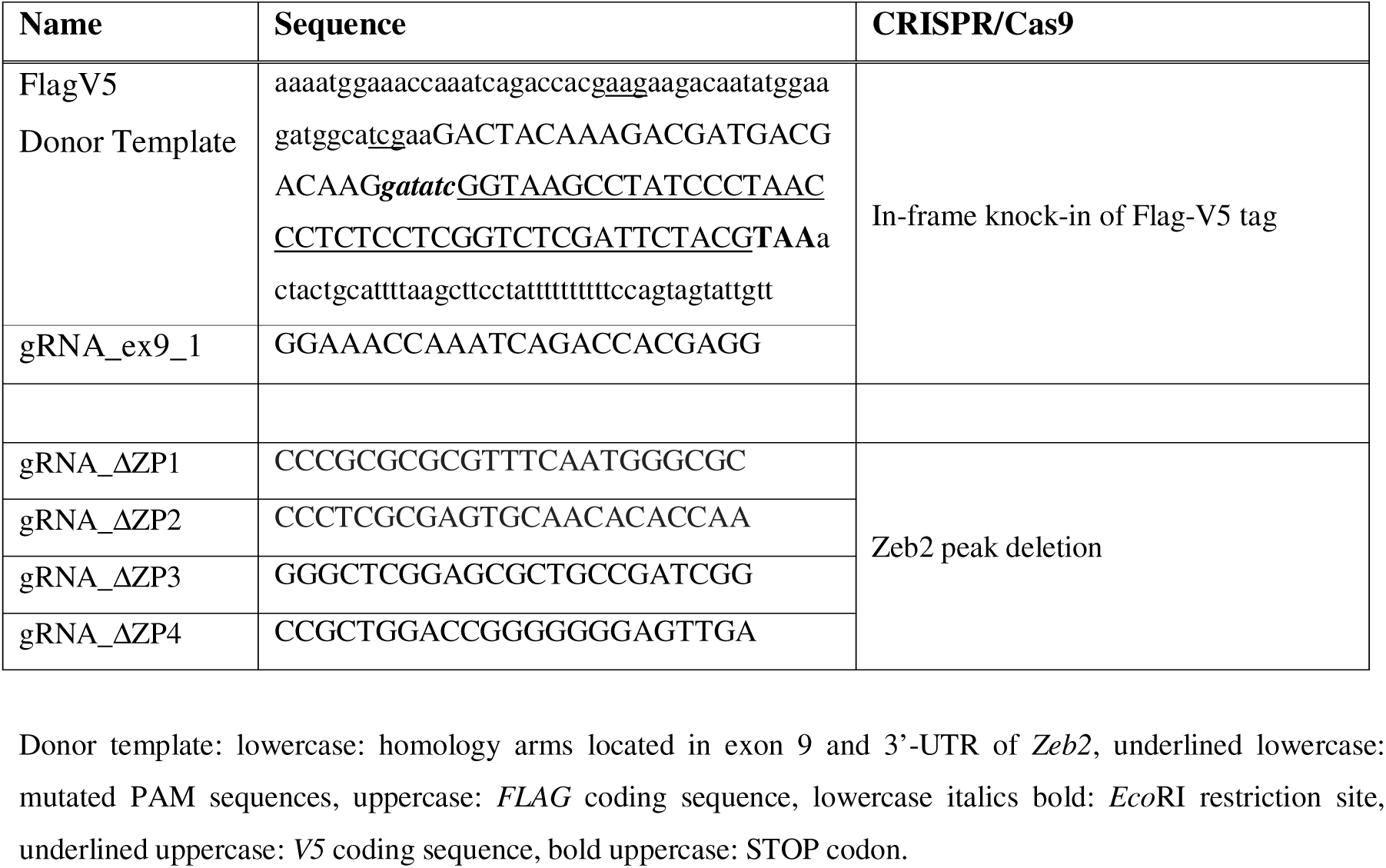
gRNAs and donor template used for CRISPR/Cas9-mediated *Zeb2* editing. Donor template: lowercase: homology arms located in exon 9 and 3’-UTR of *Zeb2*, underlined lowercase: mutated PAM sequences, uppercase: *FLAG* coding sequence, lowercase italics bold: *Eco*RI restriction site, underlined uppercase: *V5* coding sequence, bold uppercase: STOP codon.

### CRISPR/Cas9-mediated deletion of Zeb2 binding site located at chr2:45109746-45110421

Oligonucleotides for gRNAs (Table 1) with target outside of this chr2-region were cloned into *Bbs*I-digested pX330-hspCas9-T2A-eGFP plasmid. All plasmids were sequenced. 4 µg of gRNA-plasmids (1 µg each) were transfected in 350,000 mESCs and selected. After 24 hours these cells were sorted as GFP+ cells (Becton-Dickinson LSR Fortessa). Per well of a 6-well plate 1,000; 1,500; or 2,000 GFP+ cells were plated and colonies allowed to form, picked and genotyped by PCR using primers flanking this deletion, and within and outside of it. Clones showing a possible heterozygous or homozygous deletion, as concluded from the PCR analysis, were subjected to ND. At D8, they were harvested, RNA was isolated and cDNA synthesized (see below), and amplified (for the primers, see Table S1). All candidate clones were validated by Sanger-sequencing.

### ChIP-seq

DNA libraries from input (i.e. control) and V5 ChIPs were prepared using ThruPLEX DNA protocol (TakaraBio) specific for low amounts of DNA, and sequenced on Illumina HiSeq-2500, and single reads of 50 bp were generated. Adapter sequences were trimmed from the 3’-end of the reads, after which the reads were aligned to the mm10/GRCm38 genome using HISAT2 (97). From the alignments, secondary, supplementary, low quality and fragmented alignments (fragments > 150 bp) were filtered away. Peaks were called with MACS (98), and coverage was determined. 42 and 25 million reads were generated for input and V5 ChIP, respectively.

### ChIP-seq data analysis

Peak calling was performed with MACS2 (Galaxy version 2.1.1.20160309.6) (98, 99), with default parameters (narrow peak calling, *Mm*1.87e9, FDR < 0.05) using the input sample as background. The No model parameter was used, and the extension size was set on 210 bp based on the predicted fragment lengths from the alignments (MACS2 predict-tool, Galaxy version 2.1.1.20160309.1; 98, 99). The distance of the aligned reads from the TSS of the gene was analyzed using ComputeMatrix (Galaxy version 3.3.2.0.0) and PlotHeatmap Galaxy version 3.3.2.0.1; the used matrix is based on the log2ratio of the aligned ChIP peaks over the input, calculated using BamCompare (Galaxy version 3.3.2.0.0) (100).

### Transcription Factors motifs enrichment analysis

To identify the TFBS in ZEB2 binding regions associated with DEGs, we first extracted unique ZEB2 peaks located 10kb +/- from TSS. Next, we analyzed the TFBS enrichment using UniBind enrichment tool with motifs from the UniBind database (using reference genome GRCm38/mm10) (101). As a background for the analysis all ZEB2 peaks were used. The p-value from Fisher’s exact test after multitest adjustments was used to identify significantly enriched TFBS. Further, the max rank index calculated based on the odds ratio, p-value from Fisher’s exact test and the number of overlapping regions, was applied to rank the top enriched motifs.

### RNA-seq

The quality of total RNA (of biologically independent triplicates) of wild-type mESCs at D0, and at ND-D4, D6 and D8, was checked on Agilent Technologies-2100 Bioanalyzer, using an RNA nano-assay. All samples had RIN value of 9.8 or higher. Triplicate RNA-seq libraries were prepared (Illumina TruSeq stranded mRNA protocol; www.illumina.com). Briefly, 200 ng of total RNA was purified using polyT-oligo-attached magnetic beads for ending with polyA-RNA. The polyA-tailed RNA was fragmented, and cDNA synthesized (SuperScript II, random primers, in the presence of Actinomycin D). cDNA fragments were end-repaired, purified (AMPureXP beads), A-tailed using Klenow exo-enzyme and dATP. Paired-end adapters with dual index (Illumina) were ligated to the A-tailed cDNA fragments and purified (AMPureXP beads).

The resulting adapter-modified cDNAs were enriched by PCR (Phusion polymerase) as follows: 30 sec at 98°C, 15 cycles of (10 sec at 98°C, 30 sec at 60°C, 30 sec at 72°C), 5 min at 72°C. PCR products were purified (AMPureXP beads) and eluted in 30 µl resuspension buffer. One μl was loaded on an Agilent 2100 Bioanalyzer using a DNA-1000 assay to determine concentration and for quality check. Cluster generation was performed according to the Illumina TruSeq SR Rapid Cluster kit v2 Reagents Preparation Guide (www.illumina.com). After hybridization of the sequencing primer, sequencing-by-synthesis was performed using a HiSeq-2500 with a single-read 50-cycle protocol followed by dual index sequencing. Illumina adapter sequences have been trimmed off the reads, which were subsequently mapped against the GRCm38 mouse reference (using HiSat2 version 2.1.0; (97)). Gene expression values were called (using HTSeq-count version 0.9.1; 102) and Ensembl release 84 gene and transcript annotation. Sample QC and DEG analysis have been performed in the R environment for statistical computing (version 3.5.3, using DESeq2 version 1.22.1 and Tidyverse version 1.2.1 (https://github.com/tidyverse; https://www.r-project.org/; (103, R Core Team 2018 (104)).

### Meta-analysis

RNA-seq datasets (as DEG tables) were downloaded from GEO (https://www.ncbi.nlm.nih.gov/geo/, GSE35616 and GSE103003, respectively). Cross-referencing and visualization was performed in R, using Tidyverse, VennDiagram and pheatmap packages.

### Pathway-enrichment, GO, Function analysis and Gene-to-Disease association

Gene Onthology, pathway enrichment and function analysis were performed with StringDB package for R, while for Gene-to-Disease association Disgenet2R for R was used.

### Data Availability Statement

Data are available upon kind request.

## Acknowledgements

This work was supported mainly by Erasmus University Medical Center via departmental funds, extra funds coordinated with its Executive Board, and BIG project funding, a collaborative extra support between Erasmus University Rotterdam and its University Medical Center for promoting fundamental research at the Theme of Biomedical Sciences, and hence the Department of Cell Biology. We thank all co-workers of the Center for Biomics-Genomics at Erasmus University Medical Center for their expert technical assistance, and colleagues Frank Grosveld and Raymond Poot for constructive discussions.

## Legends to Supplemental Figures

**Figure S1. Characterization of the Zeb2-V5 mESC clone.**

A) Schematic overview of the design strategy, including showing also primers used for genotyping and detection of tagged allele. B) Genotyping results (selected part) showing the heterozygous band (red arrow) present in mouse (m) ESC clone 2BE3. Lower band represents the wild-type (WT) allele. C) Zeb2-V5 specific PCR showing the presence of the tagged allele only in clone 2BE3, and not in WT genomic DNA material. Clone 1BD4 was also used as negative control. D) Sanger DNA-sequencing results of clone 2BE3 showing the alignment with the Zeb2V5 sequence designed *in silico*. The last three amino acids (i.e. GME) of WT Zeb2 have been modified in DK to remove a PAM sequence and mutate the remaining PAM sequence, target of the gRNA, to avoid multiple cutting of the CAS9. An *Eco*RI restriction site was inserted between the Flag and the V5 coding sequences to facilitate for screening and an artificial STOP codon was added after the V5-coding sequence.

**Figure S2. Cross-reference of Zeb2Flagv5-bound protein-coding genes and transcriptome of differentiating mESCs.**

A) Volcano plots showing the distribution of Zeb2 bound genes in the whole transcriptomes of D4, D6 and D8 neural differentiated mESCs. Red dots depict the genes bound by Zeb2 and grey crosses those not bound. B) About 12% of the up- or down-regulated genes are bound by Zeb2 (red bars) at D8 and the percentage does not change when expanding our analysis to the other points of differentiation. C) Of the 1,952 genes bound by Zeb2 and found being differentially expressed during mESC differentiation, 335 are in common among the three considered time points. *Zeb2* itself is among these common genes and its expression increases during differentiation (yellow dot). The volcano plots show the distribution of the common genes during differentiation. D) Timepoint specific DEGs depicted as volcano plots. The grey arrow in D8 panel indicates *Tcf4*.

**Figure S3: Top 10 TF motifs enrichment at peaks present in −10/+10 kb from the TSS of up- or down-regulated genes**

Scatter plots represent the top 10 TF motifs found in the peaks present in the −10/+10 kb from the TSS range in genes which are up- or down-regulated during mESCs neural differentiation (panel A and B respectively). Each dot represent a peak.

**Figure S4. Expression of genes resulting from the meta-analysis of Zeb2-bound genes vs three independent RNA-seq datasets.**

Heatmap visualizes the log2FoldChange of the resulting genes during mESC differentiation. RNA-seq datasets where the genes have been found to be differentially expressed are annotated as green (Nkx2.1|Zeb2), purple (Zeb2 KD in mESCs) or yellow (Gsh2|Zeb2) squares.

**Figure S5. Genotyping of *Zeb2^ΔP/ΔP^* mESCs.**

A) Primers (listed in Table 1) used for genotyping. B) PCR confirming the proper deletion of the selected region in clone E9.C) Further PCR showing the difference in molecular weight for regions amplified with different set of primers and discriminating between WT, heterozygous (*Zeb2^ΔP/+^*) and homozygous (*Zeb2^ΔP/ΔP^*) deletion clones.D) Sanger sequencing showing the correct deletion of the selected region in the *Zeb2^ΔP/ΔPn^* clone.

**Figure S6. Distribution of Zeb binding sites (performed as single E-boxes; for details, see text) and TGFβ/BMP activated (phospho-)Smads and Smad4 binding elements within the identified Zeb2- bound regions for primary target genes.**

A) Grey triangles represent distance from the TSS, while blue triangles depict the TSS of the individual genes. Direction of the arrow shows also whether the genes is transcribed on the + or – strand. B) Motif analysis identifies the (half-)binding motif for Zeb2 as CACCTG.

**Figure S7. Gene-to-Disease association of the D8 DEGs bound by Zeb2.**

The gene names of the DEGs expressed at D8 of neural differentiation were initially subjected to a mouse to human gene conversion. After that, gene to disease association analysis was performed showing that a number of those genes, could be associated with neurodevelopmental disorders, mental disorders, eye defects, seizures and speech impairment.

## Supplementary Tables

**Table S1:**
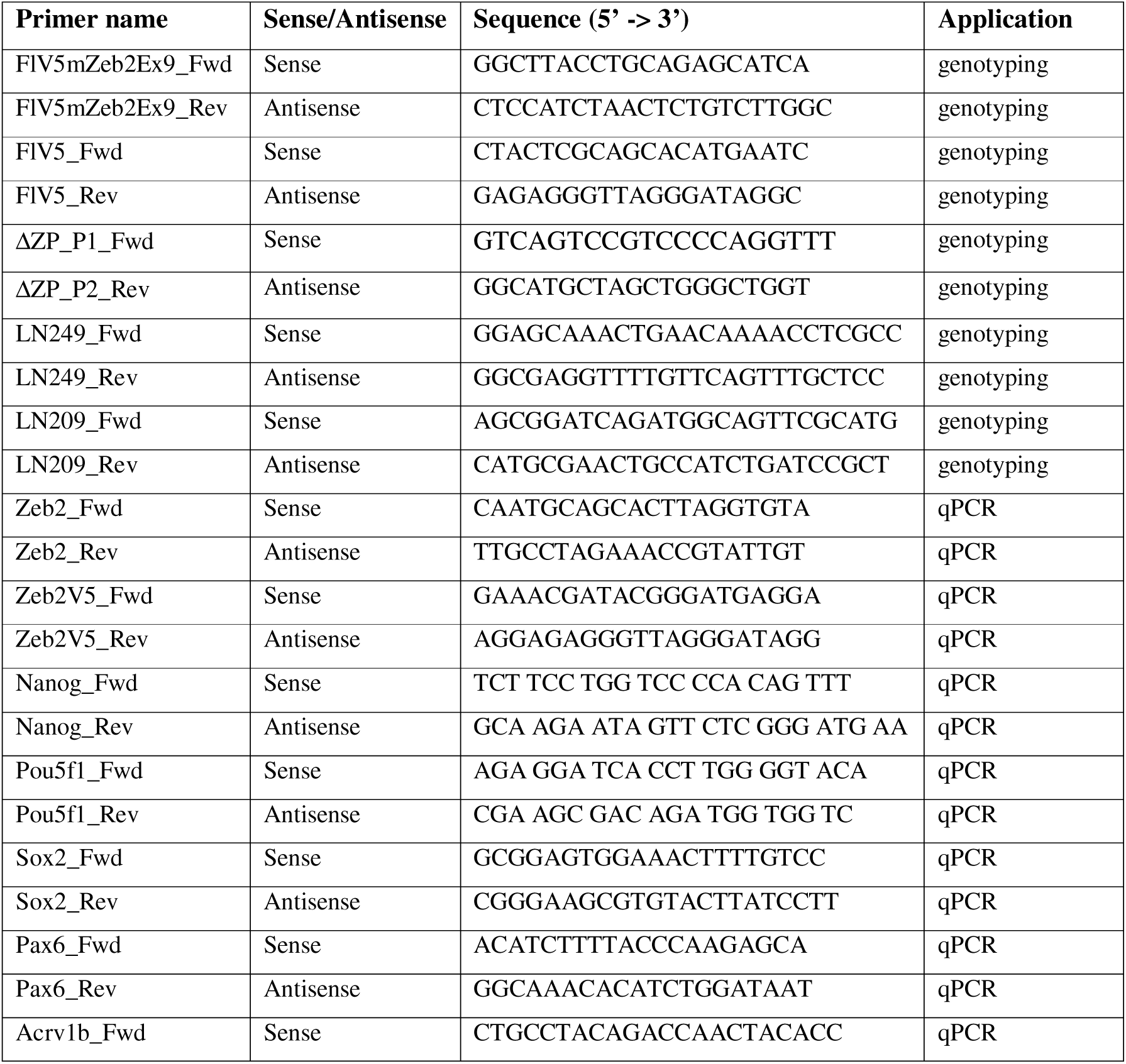
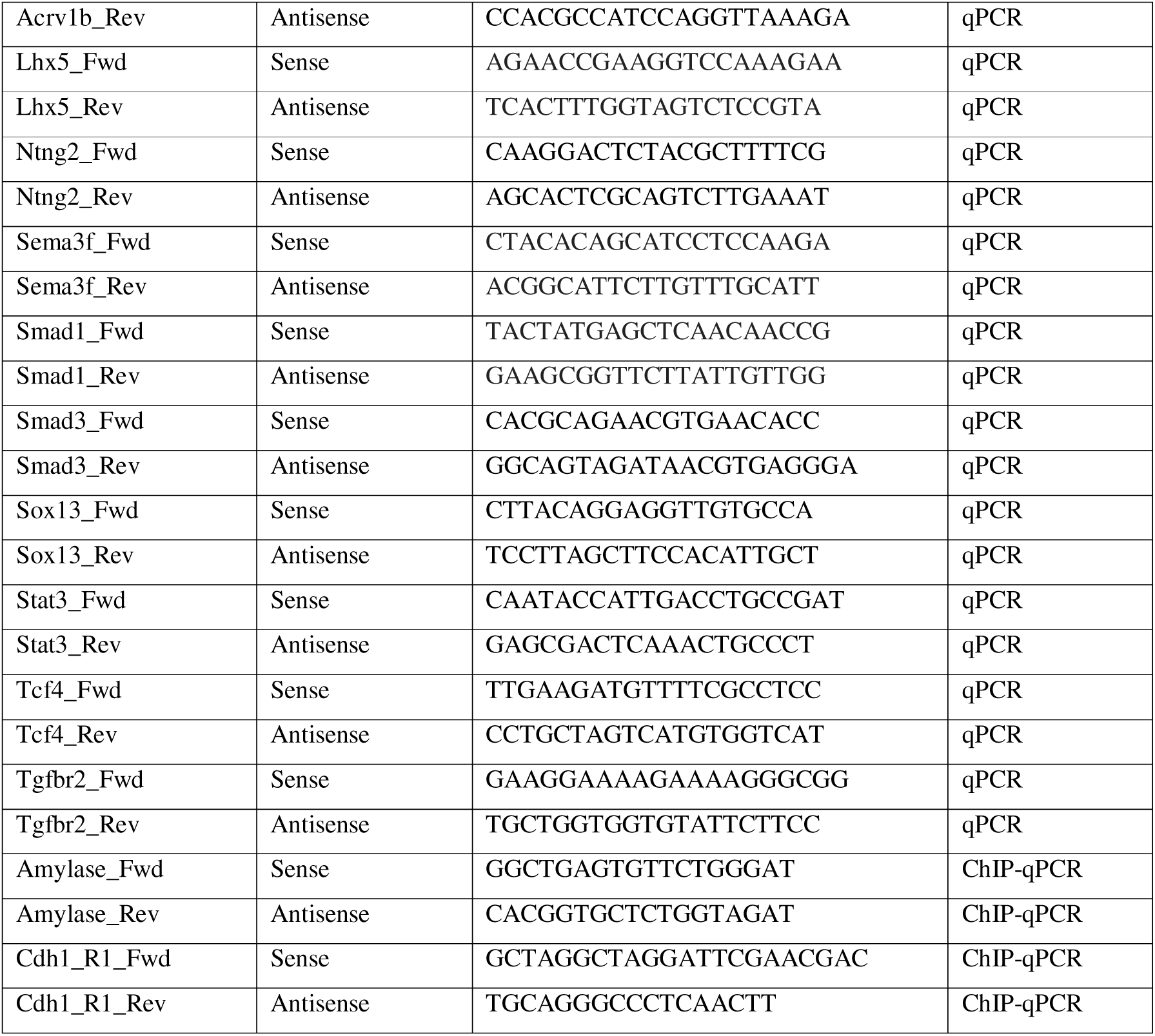
List of primers used in the study.

**Table S2:**
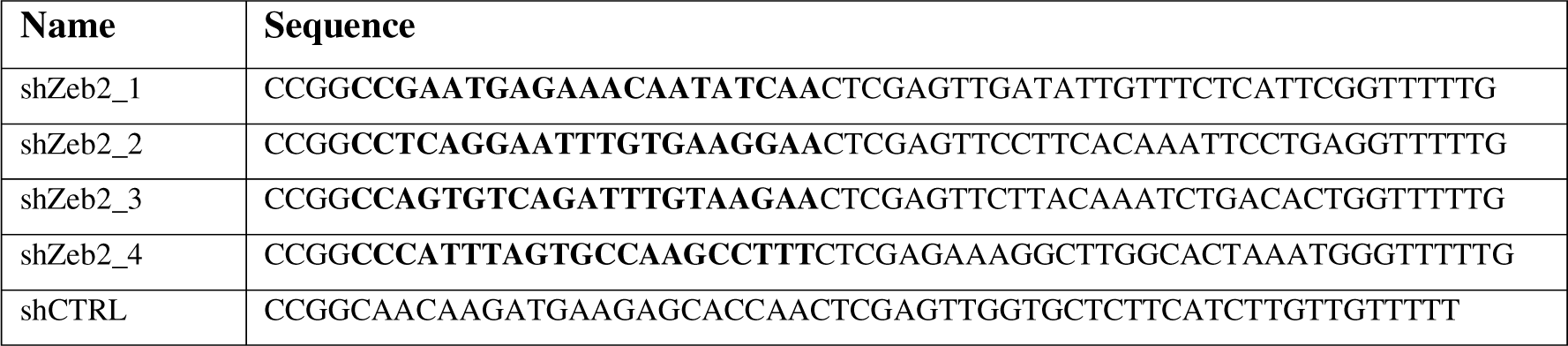
shRNAs used.

## Supplementary Files

**File S1. Narrow peaks obtained from the ChIP-seq.**

**File S2. Transcriptomic data of Zeb2 bound genes in differentiating mESCs.**

**File S3. Target genes bound by Zeb2 and deregulated in different already published RNA-seq datasets.**

## Abbreviations

bHLH: basic helix-loop-helix
BMP: bone morphogenetic protein
ChIP: chromatin immunoprecipitation
CNS: central nervous system
D: day
DEG: differentially expressed gene
EMT: epithelial-to-mesenchymal transition
EpiLSC: epiblast stem cell like cell
ESC: embryonic stem cell
esiRNA: endoribonuclease-generated interfering RNA
ex: exon
GFP: green fluorescent protein
GO: gene ontology
gRNA: guide RNA
GWBS: genome-wide binding sites
KD: knockdown
KO: knockout
lncRNA: long non-coding RNA
miR: micro-RNA
MOWS: Mowat-Wilson syndrome
ND: neurodifferentiation
NPC: neuroprogenitor\
OPC: oligodendrocyte precursor
PTHS: Pitt-Hopkins syndrome
RNA-seq: RNA-sequencing
RT: room temperature
RT-qPCR: reverse transcription quantitative polymerase chain reaction
shRNA: short-hairpin RNA
TF: transcription factor
TGFβ: transforming growth factor type β
TSS: transcription start site
V-SVZ: ventricular-subventricular zone

