## Supplementary figures and images for "Zeb2 DNA-binding sites in neuroprogenitor cells reveal autoregulation and affirm neurodevelopmental defects, including in Mowat-Wilson Syndrome"

### Supplementary Figures_R1.pdf

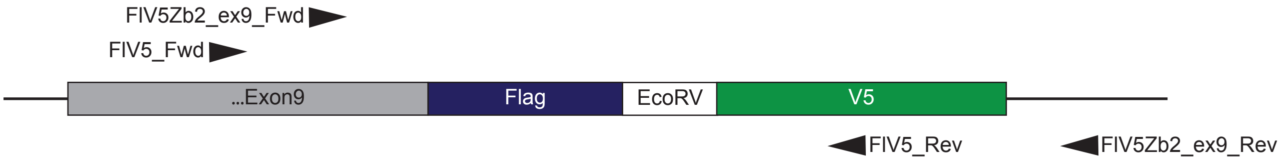

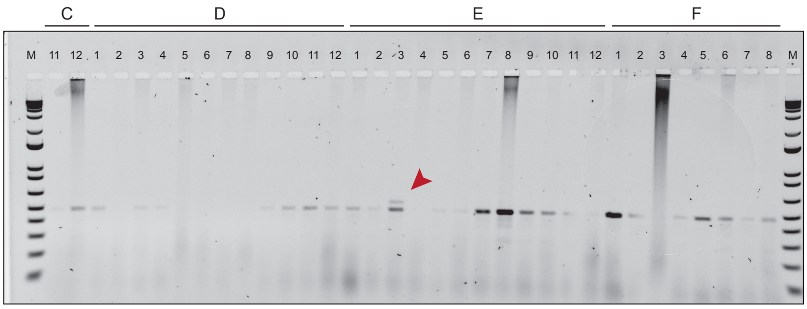

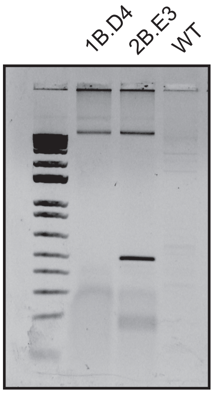

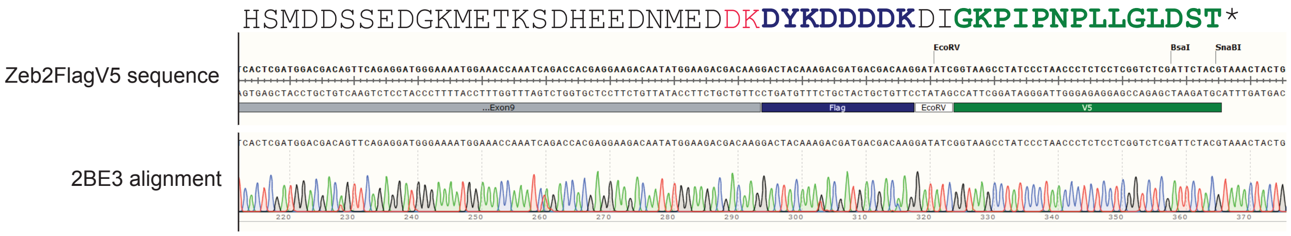

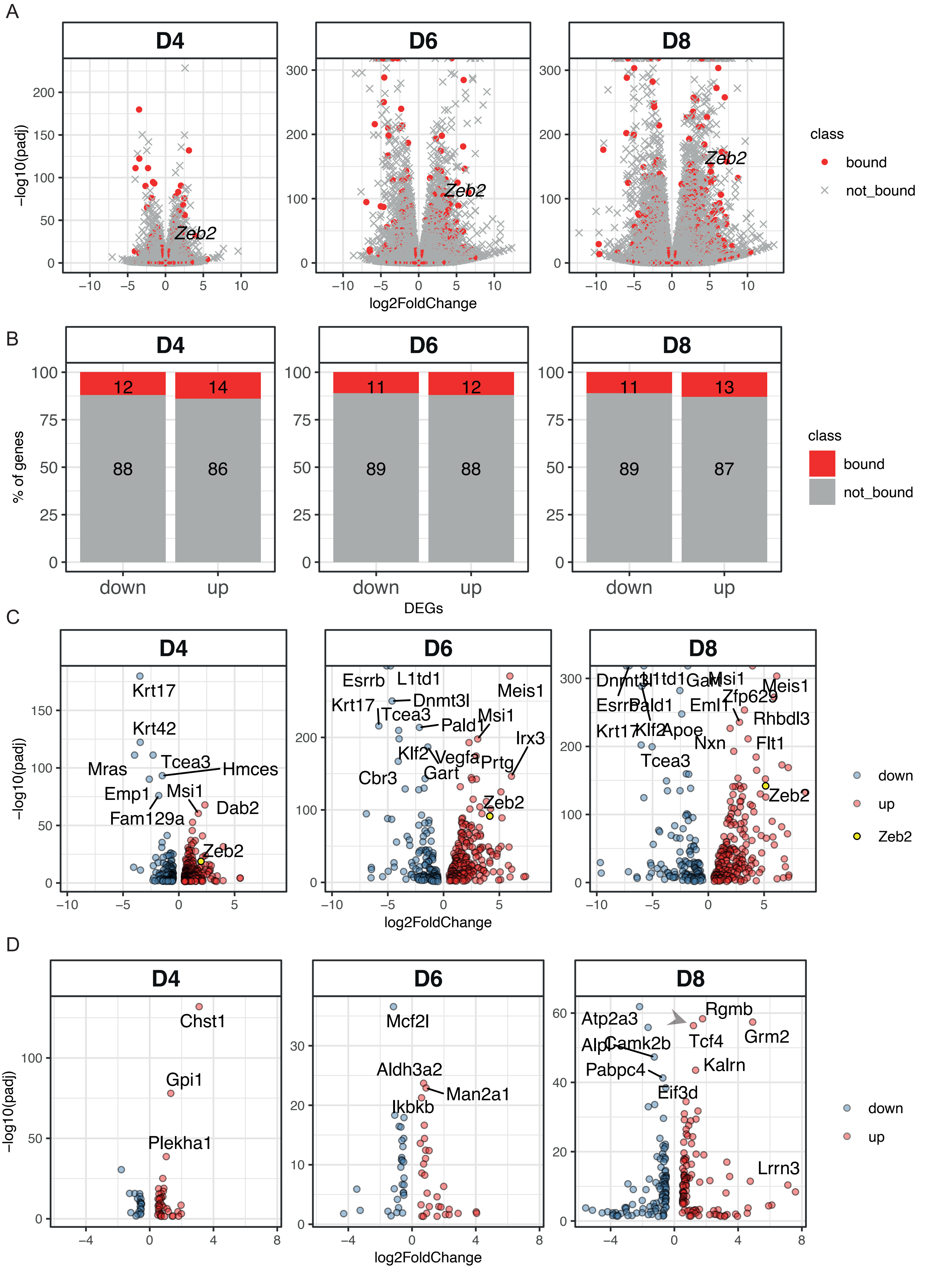

A

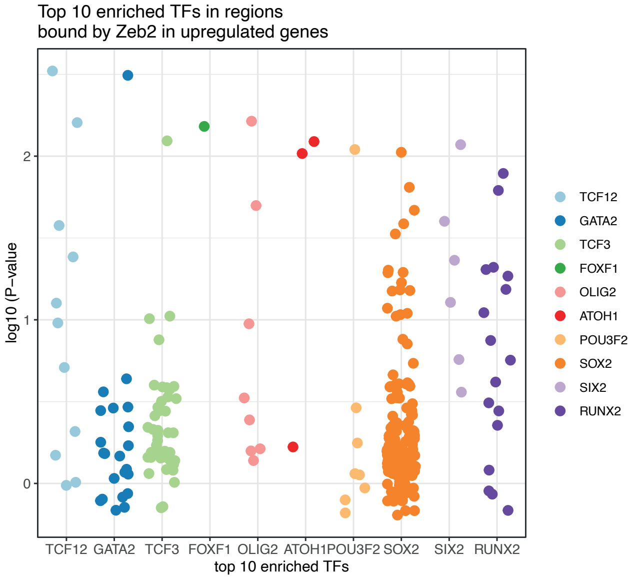


B

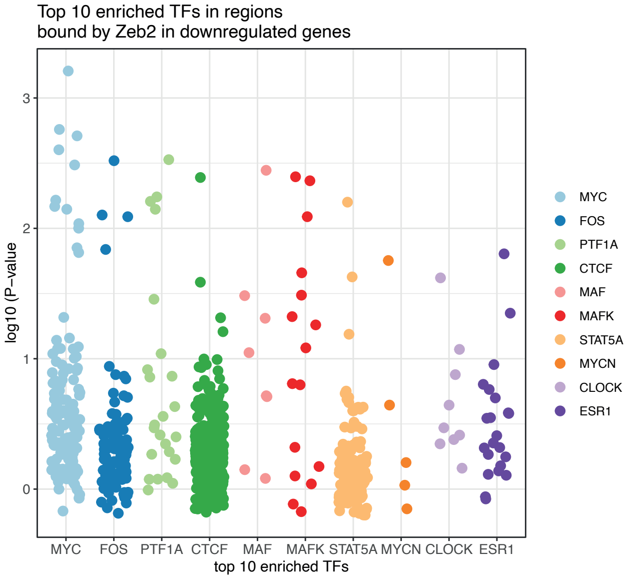


Birkhoff et al 2021_
Zeb2V5 ChIPseq

Supplementary Figure 4

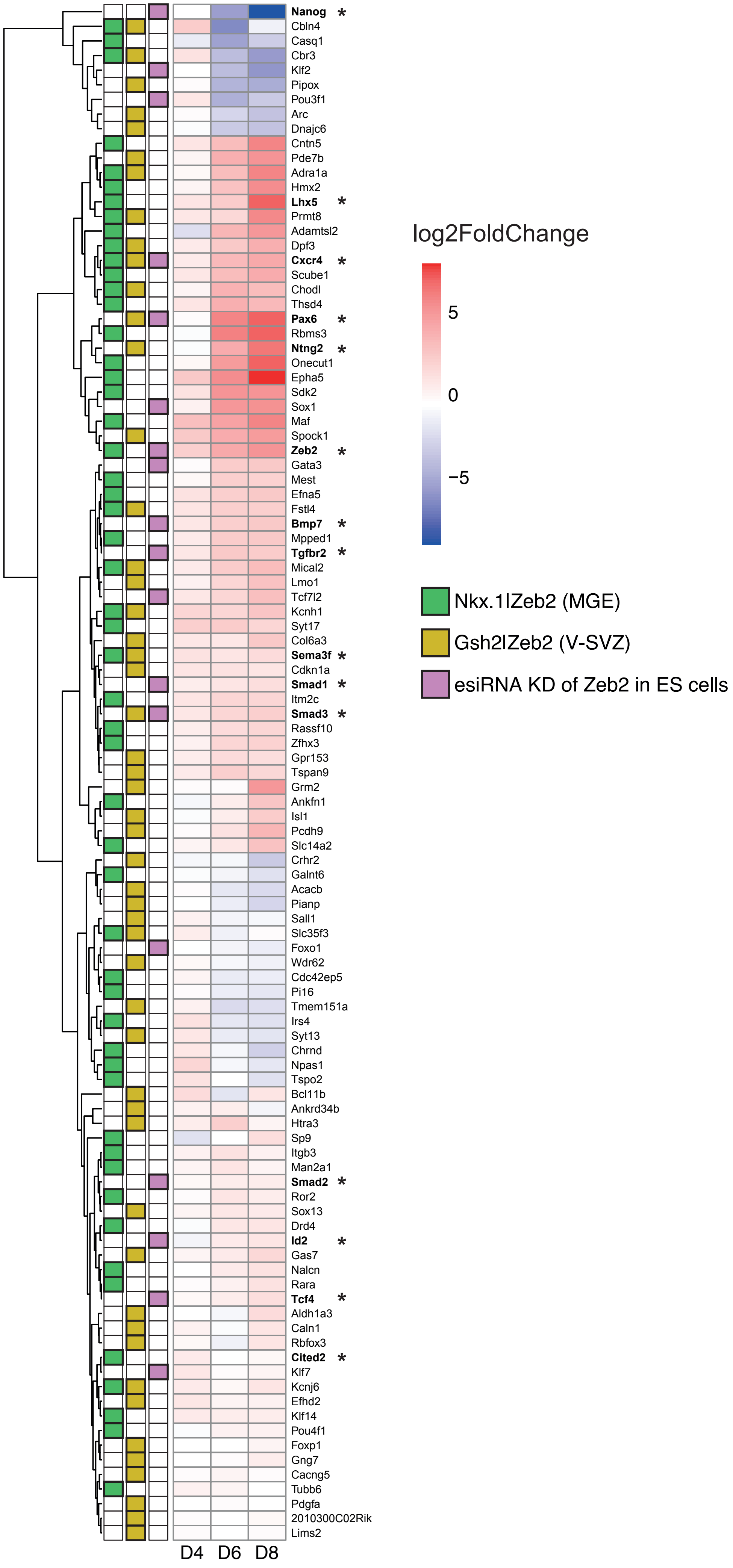

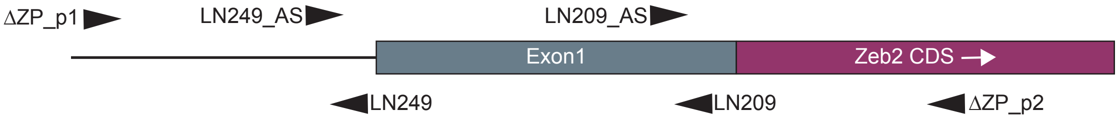

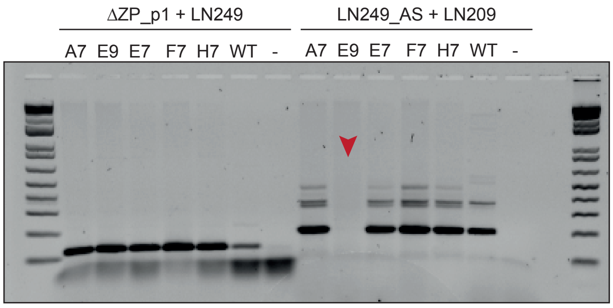

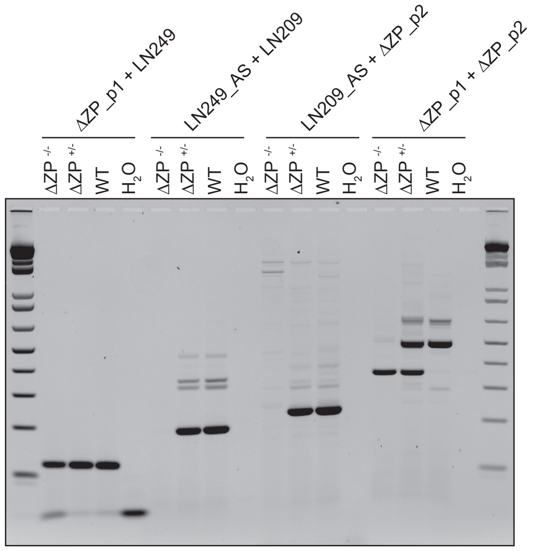

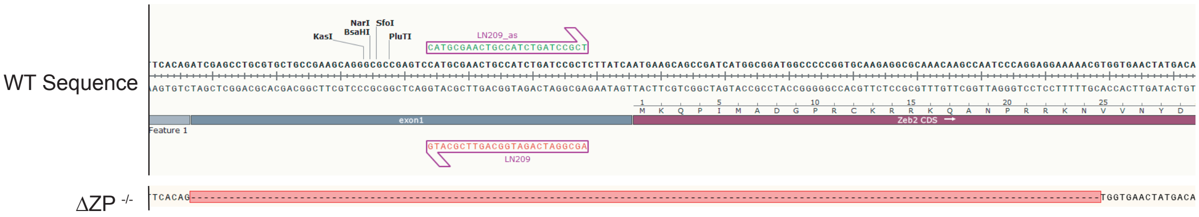

Supplementary Figure 6

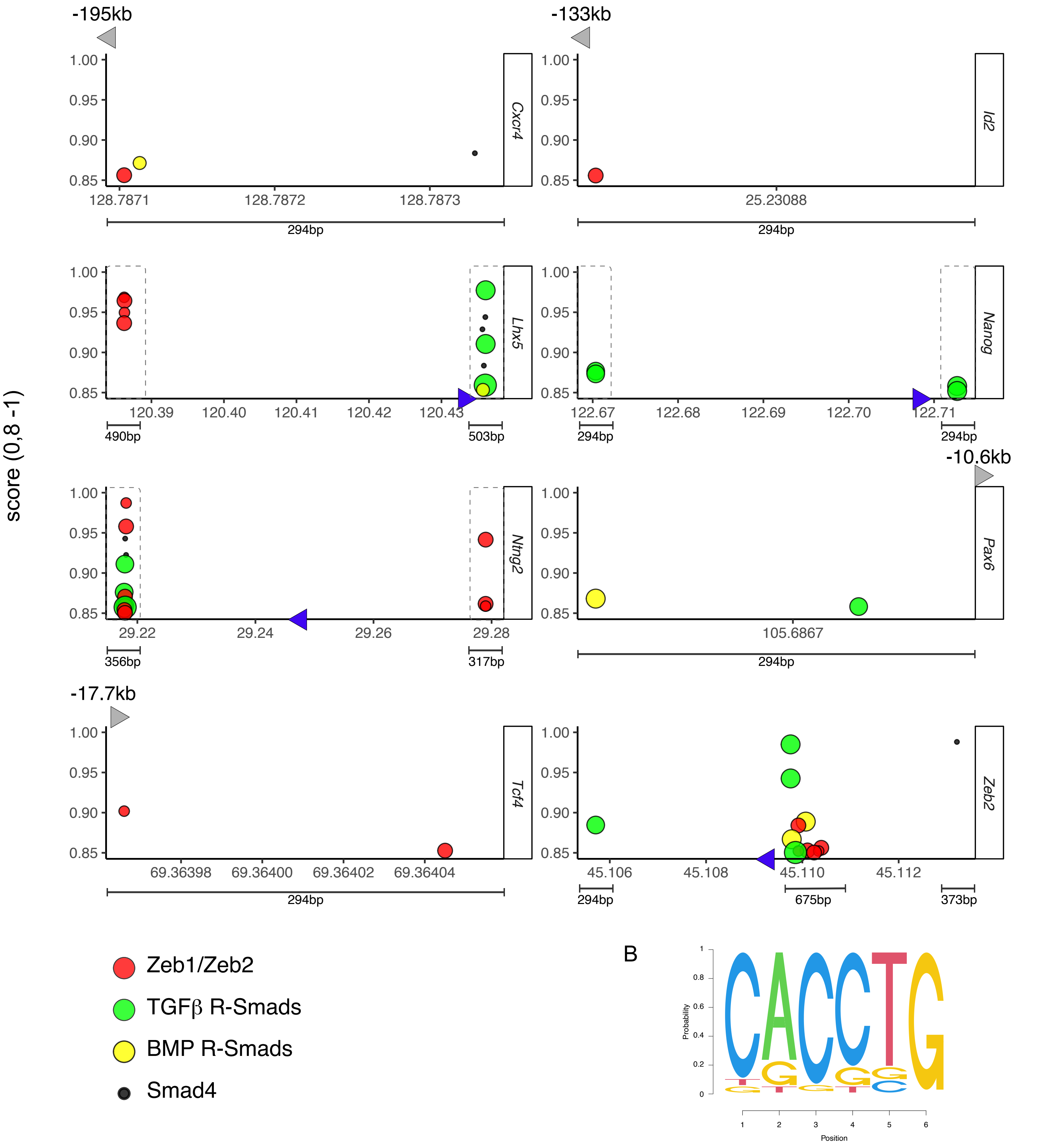

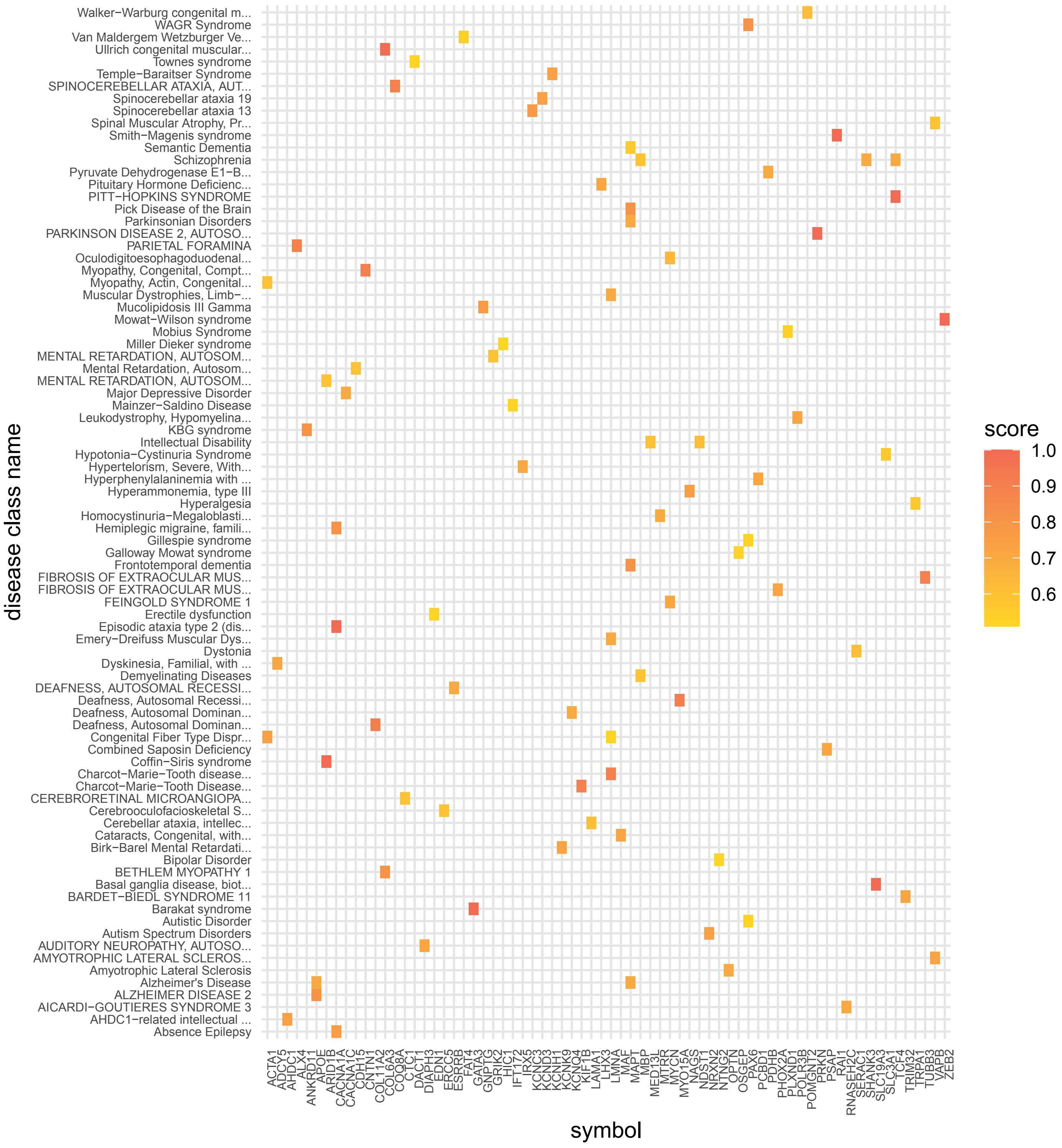
